# Reduced 4E-BP1 activity leads to acquired capivasertib resistance via increased cap-dependent protein synthesis

**DOI:** 10.1101/2025.10.10.681642

**Authors:** Helen E. Grimsley, Jasmine M. Jakubowski, Denis T. Akan, Edith Blackburn, Michelle D. Garrett

## Abstract

**Background:** The PI3K/AKT/mTOR (PAM) pathway is frequently dysregulated in human cancers and is a major target of therapeutic intervention, exemplified by the newly approved AKT inhibitor capivasertib. However, clinical responses are often limited by the emergence of acquired drug resistance.

**Methods:** To uncover potential mechanisms underlying acquired resistance to capivasertib, we developed a drug-resistant ovarian cancer model (A2780 254R) through chronic exposure to the capivasertib precursor CCT129254.

**Results:** The 254R cells exhibit broad cross-resistance to AKT inhibitors, including capivasertib, along with mTORC1 and PI3K inhibitors, in the absence of AKT mutations or elevated kinase activity. A defining feature of resistance is the reduced expression and phosphorylation of 4E-BP1, a key negative regulator of cap-dependent protein synthesis (CDPS). mcGTP pull-down assays revealed impaired 4E-BP1–eIF4E binding and persistent eIF4F complex formation, which was consistent with enhanced CDPS. An ∼9-fold increase in CDPS activity was observed in 254R cells, without corresponding increases in global protein synthesis or the expression of canonical eIF4E targets. Strikingly, ectopic expression of wild-type or constitutively active 4E-BP1 restored capivasertib sensitivity, re-established the 4E-BP1–eIF4E interaction, and significantly reduced resistance to capivasertib. These findings identify impaired 4E-BP1 function and elevated CDPS as central mediators of acquired resistance to capivasertib.

**Conclusion:** Overall, this work underscores the therapeutic vulnerability of the translational machinery to AKT inhibitor resistance. Targeting the eIF4F complex may offer a promising strategy to overcome acquired resistance, and the expression or phosphorylation of 4E-BP1 and eIF4E may serve as predictive biomarkers to guide use of capivasertib in the clinic.

## Introduction

The PI3K/AKT/mTOR (PAM) signalling axis is one of the most frequently dysregulated pathways in human malignancies^1^. Aberrant activation can occur through oncogenic mutations, amplifications, or loss of negative regulators such as PTEN, leading to enhanced cell growth, proliferation, metabolic reprogramming, and survival ^1^. AKT functions as a central signalling hub within this cascade, phosphorylating over 100 substrates that govern diverse oncogenic processes ^1^. Consequently, considerable effort has been directed toward developing pharmacologic inhibitors that target AKT, several of which have undergone clinical evaluation ^2,3^.

Capivasertib (AZD5363) is an orally bioavailable, ATP-competitive pan-AKT kinase inhibitor developed from the early lead compound CCT129254 ^4–6^. It exhibits potent and balanced inhibition across all three AKT isoforms (AKT1, AKT2, and AKT3) and has demonstrated robust anticancer activity in both *in vitro* and *in vivo* preclinical models in multiple cancer types ^4,6–11^. Capivasertib in combination with fulvestrant is the first approved AKT inhibitor for the treatment of adults with hormone receptor (HR)-positive, HER2-negative, locally advanced or metastatic breast cancer with one or more PIK3CA/AKT1/PTEN alterations ^12–17^. Despite these promising advances, as with many targeted therapies, the emergence of acquired resistance remains a significant obstacle to durable clinical responses ^18^.

Acquired resistance to AKT inhibitors typically emerges through adaptive signalling rewiring that restores critical downstream processes despite continued drug exposure ^1,19,20^. Such mechanisms may include reactivation of mTORC1 signalling or activation of alternative growth pathways ^19–21^. These changes enable tumour cells to maintain proliferative and survival signals, undermining the initial therapeutic effect. Dissecting these mechanisms is critical for improving outcomes, enabling biomarker-based patient selection, and guiding rational drug combinations.

Ovarian cancer is a malignancy in which PAM pathway activation is common, particularly in high-grade serous and endometrioid subtypes, and is linked to aggressive clinical behaviour and poor prognosis ^22–26^. Although there is a strong preclinical rationale for AKT inhibition in this setting, clinical trials have shown limited benefit, underscoring the need to identify and overcome resistance mechanisms ^27^.

Experimentally, generating and characterszing drug-resistant cell line models is a powerful strategy for elucidating acquired resistance ^28^. Sustained drug selection pressure *in vitro* can reproduce the molecular adaptations observed in patients, providing a controlled platform for detailed biochemical, genomic, and functional studies ^28,29^. Such models enable direct testing of causality, facilitate biomarker discovery, and reveal vulnerabilities that may be exploited therapeutically ^28,29^.

In this study, we generated a CCT129254-resistant derivative of the A2780 human ovarian carcinoma cell line, which exhibited marked cross-resistance to capivasertib. Mechanistically, resistance is associated with reduced phosphorylation and activation of 4E-BP1, a key negative regulator of cap-dependent translation. Functional analyses revealed that decreased 4E-BP1 activity impaired its interaction with eIF4E, thereby promoting cap-dependent protein synthesis (CDPS). Importantly, ectopic overexpression of 4E-BP1 restored sensitivity to capivasertib, functionally linking 4E-BP1 downregulation to the resistant phenotype. These findings suggest that reduced 4E-BP1 expression or activity may serve as a biomarker of acquired resistance to capivasertib and other AKT inhibitors. This work provides mechanistic insight into the translational control of the capivasertib response and identifies a potential biomarker-driven strategy for patient stratification and rational combination therapy.

## Materials and Methods

### Compounds

Compounds were sourced as follows: 4EGI-1 (Adooq, USA), capivasertib/AZD5363 (Selleck Chemicals), CCT129254 (Institute of Cancer Research, courtesy of John Caldwell and Ian Collins), cycloheximide (Sigma Aldrich), everolimus/RAD001 (Fluke Analytics), Geneticin/G418 (Santa Cruz), ipatasertib/GDC-0068 (Selleck Chemicals), MK-2206 (Selleck Chemicals), MRT00206081 (MRC Technology (LifeARC), courtesy of Andy Merritt and Ed McIver), pictilisib/ GDC-0941 (Selleck Chemicals), ravoxertinib/GDC-0994 (Selleck Chemicals), selumetinib/AZD6244 (Selleck Chemicals), and vistusertib/AZD2014 (Selleck Chemicals).

### Cell culture and transfection conditions

The A2780 human ovarian carcinoma cell line was previously purchased from the Health Protection Agency (Salisbury, UK). This cell line and subsequent drug-resistant sublines were maintained in Dulbecco’s modified Eagle’s medium high glucose supplemented with L-glutamine and 10% heat-inactivated foetal bovine serum. The cells were incubated at 37°C in a humidified 5% CO_2_ incubator. The cell lines were routinely tested approximately every 6 months to ensure that they were free from *Mycoplasma* contamination via the VenorGeM® Mycoplasma PCR detection kit (Minerva Labs, UK). Transfections with the indicated siRNAs were performed via Lipofectamine 2000 (Thermo Fisher Scientific) according to the manufacturer’s protocol.

### Generation of resistant cell lines

The A2780-derived resistant line (254R) was generated via the addition of the half-maximal growth inhibitory concentration (GI_50_) of CCT129254 to A2780 cells, which were allowed to grow until approximately 70% confluence. The cells were subsequently passaged and treated with increasing concentrations of CCT129254. This process was repeated for a period of six months.

Clonal sublines of 254R were generated via single-cell seeding. All resistant cells were routinely maintained in 56 µM CCT129254 (the maximum concentration used for generation of the resistant population) and removed from selection pressure in drug-free media one week prior to plating for experiments.

### Generation of stable overexpressing 4E-BP1 cell lines

The cells were seeded at 5x10^5^ cells in T25 flasks and allowed to adhere overnight. A total of 6.5 μg of plasmid DNA (pcDNA3, pcDNA3-4E-BP1WT-HISMYC or pcDNA3-4E-BP15A-HISMYC) was complexed with 18.75 μL of TransIT®-LT1 in OptiMEM GlutaMAX for 15c30 minutes before being added to the cells and incubated for 48 hours under normal growth conditions. The transfected cells were trypsinised, and the cells were plated across all six wells of a 6-well plate and allowed to adhere for six hours. The cells were treated with 0.25–1.5 mg/mL G418 under normal growth conditions, with the drug-treated media replenished every 2–3 days until the resistant populations developed. Single colonies were isolated once they reached a sufficient colony size (approximately 1 mm) within 11–21 days of plating. After the isolation of colonies, the cells were maintained at half the concentration of G418.

### Cell viability assay

Cells were seeded into 96-well plates at the indicated concentrations in 160 μl of complete DMEM and allowed to attach for 48 hours prior to treatment. To investigate cross-resistance to different compounds, cells were treated with a range of concentrations of serially diluted compounds (1.5-5-fold) in complete DMEM, and 40 μl of each concentration was added to each well and incubated for 96 hours. Cells were then fixed with 10% (w/v) trichloroacetic acid for 30 minutes and washed three times with water. Fixed cells were stained with 0.4% (w/v) sulforhodamine B (SRB) solubilised in 1% (v/v) acetic acid for 30 minutes and washed with 1% (v/v) acetic acid. The plates were dried at 37°C. SRB was solubilised with 100 μl of 10 mM Tris-base per well, and absorbances were read at a wavelength of 490 nm in a Victor X4 multilabel plate reader (PerkinElmer Life Sciences, USA). The raw values were blanked using cell-free wells with 10 mM Tris and normalised to a percentage of the average of the untreated control per drug. The GI_50_ values were determined via GraphPad Prism 6 (GraphPad Software Inc., USA). The resistance factor (RF) values were calculated as the ratio of the GI_50_ of CCT129254-resistant cells to the A2780 GI_50_.

### AKT *in vitro* kinase assay

Cells were plated at a density of 2x10^6^ per 15 cm dish and allowed to adhere for 60 hours. The cells were washed twice with ice-cold PBS and incubated for 5 minutes in 750 μL of nondenaturing lysis buffer (1X Cell Lysis Buffer (20 mM Tris-HCl pH 7.5, 150 mM NaCl, 1 mM disodium EDTA, 1 mM EGTA, 1% Triton, 2.5 mM sodium pyrophosphate, 1 mM sodium orthovanadate, 1 μg ml-1 leupeptin; Cell Signaling Technology, USA), 10 mM β-glycerophosphate, and 1 mM phenylmethanesulfonyl fluoride). Cells were scraped off the dish and collected in microcentrifuge tubes and incubated on ice for 30 minutes. The lysates were then cleared by centrifugation at 14,000 x g for 10 minutes at 4°C. The protein concentration was estimated via BCA assay following manufacturer’s instructions, and samples diluted with nondenaturing lysis buffer to a concentration of 2.5 μg/L. To isolate AKT from cell lysates, immunoprecipitation was performed. A total of 1250 μg of lysate was precleared with 50 μL of washed Protein A agarose beads (100 μL of 50% bead mixture; Pierce Biotechnology, USA) by incubation on a rotating wheel for 30 minutes at 4°C. The samples were subsequently centrifuged at 5,000 xg for three minutes, after which the precleared lysate was collected. To bind the antibody to AKT, 500 μg of precleared lysate was incubated either as an antibody negative control or with 8 μL of AKT antibody (Cat# 4691; Cell Signaling Technology, USA) on a rotating wheel for two hours at 4°C. The samples were subsequently incubated with 20 μL of washed Protein A agarose beads on a rotating wheel for two hours at 4°C. The AKT-bound bead complexes were centrifuged at 5,000 xg for three minutes and washed twice with nondenaturing lysis buffer at 4 °C. The AKT-bound bead complexes were washed twice with 1x kinase buffer (25 mM Tris pH 7.5, 5 mM β-glycerophosphate, 2 mM DTT, 0.1 mM sodium orthovanadate, and 10 mM MgCl_2_; Cell Signaling Technology, USA). Kinase buffer (50 μL) containing 1 μg of the GSK-3 fusion protein and 200 μM ATP (Cell Signaling Technology, USA) was added to the isolated AKT, which was subsequently incubated with gentle shaking for 30 minutes at 30°C. To stop the reaction, 25 μL of 3x loading buffer (187.5 mM Tris pH 6.8, 6% (w/v) SDS, 30% glycerol, 15% (v/v) β-mercaptoethanol, 0.15% (w/v) bromophenol blue)) was added, and the samples were incubated at 95°C for five minutes. A 15 μL protein sample was subsequently analysed via western blotting.

#### Antibodies

4E-BP1 (Cat# 9644, Cell Signaling Technology), 4E-BP1 pT37/T46 (Cat#2855, Cell Signaling Technology), AKT (Cat# 4691, Cell Signaling Technology), AKT pS473 (Cat# 9271, Cell Signaling Technology), AKT pT308 (Cat# 4056, Cell Signaling Technology), cleaved PARP (Cat# 9541, Cell Signaling Technology), c-Myc (G-4) (sc-377552, Santa Cruz), Cyclin D1 (HD-11) (SC-246, Santa Cruz), eIF4E (Cat# 2067, Cell Signaling Technology), eIF4G (Cat# 2469, Cell Signaling Technology), ELK (sc-365876, Santa Cruz), ELK pS383 (sc-8406, Santa Cruz), ERK (Cat# 4695, Cell Signaling Technology), ERK pT202/Y204 (Cat# 4370, Cell Signaling Technology), GAPDH (MAB374 Chemicon), GSK-3β (Cat# 9315, Cell Signaling Technology), GSK-3β pS9 (Cat# 5558, Cell Signaling Technology), His-probe Antibody (H-3)

### Sequencing of AKT isoforms

A total of 5 × 10^6^ cells were pelleted by centrifugation at 2,000 rpm for two minutes at 4°C. The pellets were washed with ice-cold PBS. RNA was extracted via the QIAGEN RNeasy Mini Kit (QIAGEN, Germany) following the manufacturer’s instructions. Briefly, the cell pellets were resuspended in RLT buffer and homogenied by centrifugation through a QIAshredder spin column for two minutes at 14,000 x g.

RNA was precipitated from the homogenised lysate by the addition of one volume of 70% ethanol, transferred to an RNeasy spin column, and centrifuged for 15 seconds at 8,000 × *g*. Bound RNA was washed by the addition of Buffer RW1 and further centrifuged for 15 seconds at 8,000 × *g*. The column was washed again with Buffer RPE and centrifuged for 15 seconds at 8,000 × *g* and subsequently washed with Buffer RPE and centrifuged for two minutes at 8000 × *g.* A further centrifugation for one minute at 8,000 × *g* was performed. The RNA was eluted from the column with 50 μL of RNase-free water and centrifuged at 8,000 × *g* for one minute. DNA digestion was performed for 15 minutes at 37°C with 10 units of Roche DNase I recombinant, RNase-free (Roche Diagnostics GmbH, Germany), which was inactivated by the addition of mM EDTA (pH 8.0) and incubation at 75°C for 10 minutes. RNA was subsequently precipitated by the addition of ethanol to a final concentration of 35%, purified by binding to an RNeasy spin column, washed twice with Buffer RPE, and eluted following the procedure described above. The RNA concentration was determined via a NanoDrop 1000 Spectrophotometer (Thermo Scientific, USA). To reverse transcribe RNA to cDNA, Superscript II Reverse Transcriptase (Invitrogen, USA) was used. Briefly, 5 μg of purified RNA was mixed with 500 μM dNTP mixture (Invitrogen, USA) and 2.5 μM random hexamer primers (Applied Biosystems, USA) and incubated at 65°C for five minutes. First, strand buffer and 0.1 M DTT were added, and the mixture was incubated at 25°C for two minutes. Superscript II reverse transcriptase was added, and first-strand cDNA synthesis was performed by incubation at 25°C for 10 minutes, followed by incubation for 50 minutes at 42°C and subsequent incubation for 15 minutes at 70°C. To digest the RNA and leave single-stranded cDNA, RNase H was added, and the mixture was incubated at 37°C for 20 minutes. PCR was performed using cDNA as a template and Platinum® *Taq* DNA Polymerase, High Fidelity (Invitrogen, USA). A 50 μL reaction mixture containing 2 μL of cDNA and 1 unit of *Taq* Polymerase in 1x high-fidelity buffer, 2 mM MgSO4, 0.2 mM dNTP mixture, 0.2 μM forward primer and 0.2 μM reverse primer was used. The PCR cycling conditions were 40 cycles of denaturation at 94°C for 30 seconds (one minute for the first cycle), annealing at 50°C for 30 seconds, and elongation at 68°C for 150 seconds. The PCR products were purified via a QIAGEN PCR Purification Kit (QIAGEN, Germany) following the manufacturer’s instructions. Five volumes of Buffer PB were added to one volume of PCR sample and transferred to a QIAquick spin column. The DNA was bound to the column by centrifugation at 14,000 xg for 60 seconds. The DNA was washed with Buffer PE and centrifuged at 14,000 xg for 60 seconds, after which it was further centrifuged for one minute to remove residual buffer. DNA was eluted in 50 μL of TE buffer and quantified via a NanoDrop 1000 Spectrophotometer (Thermo Scientific, USA). Sequencing of premixed DNA and primer was performed via the BigDye Terminator v3.1 Cycle Sequencing Kit with a 3730xl DNA Analyser (both Applied Biosystems, USA) by Eurofins Genomics (Ebersberg, Germany). The primers used for PCR and sequencing were as follows (5’-3’):

AKT1: F1 (GCCAGCCTGGGTCAAAGAA), R1 (CACCCGGAGAACAAACTGGA), F2 (CCATGTGGAGACTCCTGAGG), R2 (CCTTCTTGAGCAGCCCTGAA), F3 (CTGCACTCGGAGAAGAACGT), R3 (GGTCGTGGGTCTGGAAAGAG), F4 (TCGGAGACTGACACCAGGTA), and R4 (GCGCCACAGAGAAGTTGTTG). AKT2: F1 (CTGTCCACGGTGCCTCCT), R1 (AAATGCGAGTCTGGGCACAA), F2 (GACGAGAGGGAGGAGTGGAT), R2 (TCCTTCTTAAGCAGCCCAGC), F3 (TTGAGTACTTGCACTCGCGG), R3 (AAGGCATACTTCAGCGCAGT), F4 (CCGAGGTCGACACAAGGTAC), and R4 (TCAGCTGGCATTCTGCTACG). AKT3: F1 (CCAAACCCTAAAGCTGATATCACA), R1 (AGCAGCAACAGCATGAGACC), F2 (TGGACAGAAGCTATCCAGGC), R2 (GTCCTCCACCAAGGCGTTTA), F3 (ACCGCACACGTTTCTATGGT), R3 (ACGGTCTTTTGTCTGGAAGGA), F4 (GCTTGTACCTCCTTTTAAACCTCA), and R4 (GCCACTGAAAAGTTGTTGAGGG).

### Immunoblotting

Cells were scraped into ice-cold lysis buffer (50 mM HEPES, pH 7.4), 250 mM NaCl, 0.1% Nonidet-P40, 1 mM DTT, 1 mM EDTA, pH 8.0, 1 mM NaF, 10 mM β-glycerophosphate, 0.1 mM sodium orthovanadate and Complete™ protease inhibitor cocktail (Roche, Switzerland). The lysates were incubated on ice for 30 minutes before centrifugation at 14,000 × g for 10 minutes at 4°C. The lysate was kept on ice for immediate use or snap-frozen on dry ice and stored at -80°C. The protein concentration was determined via the bicinchoninic acid (BCA) assay according to the manufacturer’s instructions (Thermo Fisher). Cell lysates were normalised and diluted in 5X sample buffer (312.5 mM Tris pH 6.8, 6% (w/v) SDS, 50% glycerol, 25% (v/v) β-mercaptoethanol, 0.25% (w/v) bromophenol blue) before denaturation at 95°C for five minutes. Protein samples were loaded into 6%, 10% or 15% Tris-glycine gels (6–15% acrylamide/bis(v/v), 0.375 M Tris pH 8.8, 0.1% SDS (w/v), 0.05% ammonium persulfate (w/v) and 0.005% tetramethylethylenediamine) with a 4% tris-glycine stacking gel (4% acrylamide/bis(v/v), 0.125 M Tris pH 6.8, 0.1% SDS (w/v), 0.05% ammonium persulfate (w/v) and 0.01% tetramethylethylenediamine). The gels were run in Tris-glycine running buffer (25 mM Tris, 190 mM glycine, 0.1% SDS) at 150 V for 60c90 minutes. Dual precision-prestained protein markers (Bio-Rad, USA) or Page Ruler Plus prestained protein ladders (Thermo Fisher Scientific, USA) were used as protein markers. The proteins were transferred to methanol-activated 0.2 μm pore Immobilon-P PVDF membranes (Millipore, USA) via a wet transfer system at 100 V for 90–120 minutes in prechilled transfer buffer (25 mM Tris-base, 190 mM glycine, and 10% (v/v) methanol). To investigate very high-molecular-weight proteins (i.e., eIF4G, 220 kDa), resolved 6% gels were transferred overnight (18–24 hours) at 30 V in a cold room at 4°C. The membranes were blocked on a rocking platform at RT for one hour in the following blocking buffer: Tris-buffered saline (TBST; 50 mM Tris pH 8.0, 150 mM NaCl, 0.1% Tween-20 (v/v)) containing 5% milk powder. The membranes were subsequently incubated with primary antibodies diluted in blocking buffer overnight at 4°C. The membranes were washed twice for 10 minutes with TBST and incubated with a goat anti-rabbit or anti-mouse secondary horseradish peroxidase-conjugated (HRP) antibody (Bio-Rad, USA; Table 2.1) for one hour at RT. The membranes were washed four times for five minutes each, and detection was performed via Enhanced Chemiluminescence (ECL) Western Blotting Substrate (Pierce Biotechnology, USA) or Clarity™ Western ECL Blotting Substrates (Bio-Rad, USA). Bands were developed by exposure to Amersham Hyperfilm ECL (GE Healthcare, UK) or developed via the Syngene GBox system and Genesys software (Syngene, USA). The membranes were probed for phosphorylated proteins and stripped for detection of total protein expression. The antibodies were stripped from the membrane by incubation in stripping buffer (50 mM glycine, 1% (w/v) SDS, pH 2.0) for five minutes on a rocking platform, washed twice in TBST for five minutes each, and re-probed with relevant antibodies overnight at 4°C or 1–3 hours at RT, depending on the quality of the antibody. The band density was quantified via ImageJ software. The images were imported, and the band density was calculated as a percentage of the sum density of the compared analysed bands. Each sample band was normalised to its respective loading control.

### Coimmunoprecipitation

Treated cells were lysed via 200 μL of nondenaturing lysis buffer (NDLB; 1X Cell Lysis Buffer: 20 mM Tris-HCl pH 7.5, 150 mM NaCl, 1 mM disodium EDTA, 1 mM EGTA, 1% Triton, 2.5 mM sodium pyrophosphate, 1 mM sodium orthovanadate and 1 μg ml-1 leupeptin; Cell Signaling Technology, USA), lysates prepared and protein concentration determined for the AKT kinase assay. The lysates were diluted to 0.5 mg/mL in NDLB. Fifteen microliters of immobilised γ-aminophenyl-m7GTP agarose beads (30 μL of 50% slurry; Jena Bioscience, Germany) were washed twice with 200 μL of NDLB. A 250 μg sample of lysate was added to the washed beads and incubated on a rotating wheel at 4°C overnight. The beads were centrifuged at 2,500 × g for one minute to separate the output lysate from the beads, and the beads were then washed three times with 400 μL of NDLB. Then, 50 μL of 2X sample buffer was added, and the bound proteins were denatured by incubating the beads at 60°C for 10 minutes to prevent agarose melting. Bead-free input and output lysate samples were prepared. eIF4G and 4E-BP1 levels bound to eIF4E in prepared bead samples were analysed by western blotting. eIF4E was used as an equivalent loading control between sample conditions.

### Dual-luciferase reporter assay

Cells were plated at 1x10^5^ cells per well in six-well plates and allowed to adhere overnight. The pRL-IRES-FL plasmid, pcDNA5-FRT-eGFP (both at 2.5µg) or RNase-free ddH2O were complexed with TransIT®-LT1 (at 7.5µg) and added to the cells as per manufacturers instructions. The cells were incubated under normal growth conditions for 48 hours post-transfection to allow gene expression before lysis. Lysis and determination of luciferase expression were carried out according to the manufacturer’s instructions from the Dual-Luciferase Reporter Assay System (Promega, USA). Briefly, the protein concentration of the cleared lysates was determined via BCA assay according to the manufacturer’s instructions, and the lysates were normalised to the same protein concentration in 1X Passive Lysis Buffer. To determine firefly luciferase levels, 100 μL of Luciferase Assay Reagent II was added. Luciferase activity as light emitted was immediately measured via the default luciferase protocol on a Victor X4 multilabel plate reader (PerkinElmer Life Sciences, USA), utilising a one second measurement period. The firefly luciferase reaction was stopped, and the *Renilla* luciferase reaction was initiated by adding 100 μL of Stop & Glo reagent and mixing thoroughly by pipetting. *Renilla* luciferase activity was immediately measured as described above.

### Polysome profiling

Cells were plated at 15x10^6^ in T175 flasks and allowed to adhere overnight. At the time of harvest, the cells were spiked with cycloheximide at a final concentration of 0.1 mg/mL and incubated for five minutes at 37°C. The culture medium was removed, and the cells were washed twice with ice-cold 0.1 mg/mL cycloheximide in PBS. Ice-cold polysome profiling lysis buffer (300 mM NaCl, 15 mM Tris-HCl pH 7.5, 15 mM MgCl_2_, 0.5% (v/v) Triton X-100, 5 mM β-mercaptoethanol, 0.5 mM phenylmethylsulfonyl fluoride (PMSF)) and 0.1 mg/mL cycloheximide in RNase/DNase-free H_2_O) were added, and the cells were scraped and lysates collected in prechilled RNase/DNase-free microcentrifuge tubes and incubated on ice for five minutes. Lysates were cleared of insoluble material by centrifugation at 10,000 × g for three minutes at 4°C, transferred to a fresh RNase/DNase-free microcentrifuge tube and kept on ice for immediate use or snap-frozen on dry ice for storage at -80°C. The protein concentration of the lysate was determined via the BCA assay according to the manufacturer’s instructions (Thermo Fisher). Lysate (1 mL) was loaded onto separate 10–50% RNase/DNase-free sucrose gradients (Acros Organics, USA) in Beckman 14x96 mm polyallomer tubes (Beckman Coulter, USA) and centrifuged at 180,000 × g for 135 minutes at 4°C. The absorbance of the gradient fractions was read at a wavelength of 254 nm in a BR-188 Gradient Fractionation System (Brandel, USA) to draw a physical or digital trace over time. Traces were analysed either by importing the digital copy into Microsoft Excel 2016 software or by scanning the hard copy via CanoScan LiDE120 (Canon., Japan), tracing in Inkscape and calculating the area under the curve via ImageJ software ^30^. The areas under the curves were calculated separately to measure the relative subpolysome and polysome percentages.

### Data analysis and statistics

Statistical analysis for the various experiments was performed via GraphPad Prism version 6. The results are presented as the means ± standard errors of the means (SEMs). A p value of <0.05 according to Student’s t test was considered statistically significant. Ns, nonsignificant, *p<0.05, **p<0.005, ***p<0.001.

## Results

### mTORC1 drives capivasertib resistance via a distinct role outside of the AKT/mTORC1 axis

The A2780 human ovarian cancer cell line was selected for these studies based upon its predicted sensitivity to AKT inhibition, owing to genetic alterations that reduce PTEN expression and increase PI3K activity ^31,32^. To model acquired resistance, an A2780-derived resistant line (254R) was generated via dose escalation with the early capivasertib analogue CCT129254. Compared with A2780 cells, the 254R-treated subline revealed a 23.2-fold increase in resistance (resistance factor = A2780 GI₅₀/254R GI₅₀; **Fig. 1A, Sup. Fig. 1A**). Notably, 254R showed even greater resistance to capivasertib and ipatasertib, with ∼100-fold and ∼50-fold resistance, respectively, underscoring its utility as a model of acquired resistance to ATP-competitive AKT inhibitors (**Fig. 1A, Sup. Fig. 1A**). Cross-resistance was also observed with MK-2206, a noncompetitive allosteric AKT inhibitor, with a resistance factor of 27.2-fold, suggesting that resistance at 254R is not due to alterations at the AKT binding site (**Fig. 1A, Sup. Fig. 1A**). This finding was further supported *in vitro* kinase assays and western blot analysis, which revealed no increase in AKT enzymatic activity or AKT expression respectively (**Sup. Fig. 1B, 1C**). Furthermore, sequencing revealed there were no mutations in two AKT isoforms in 254R (**Sup. Fig.1D**). We were unable to successfully amplify AKT3 in either of the cell lines.

**Figure 1:**
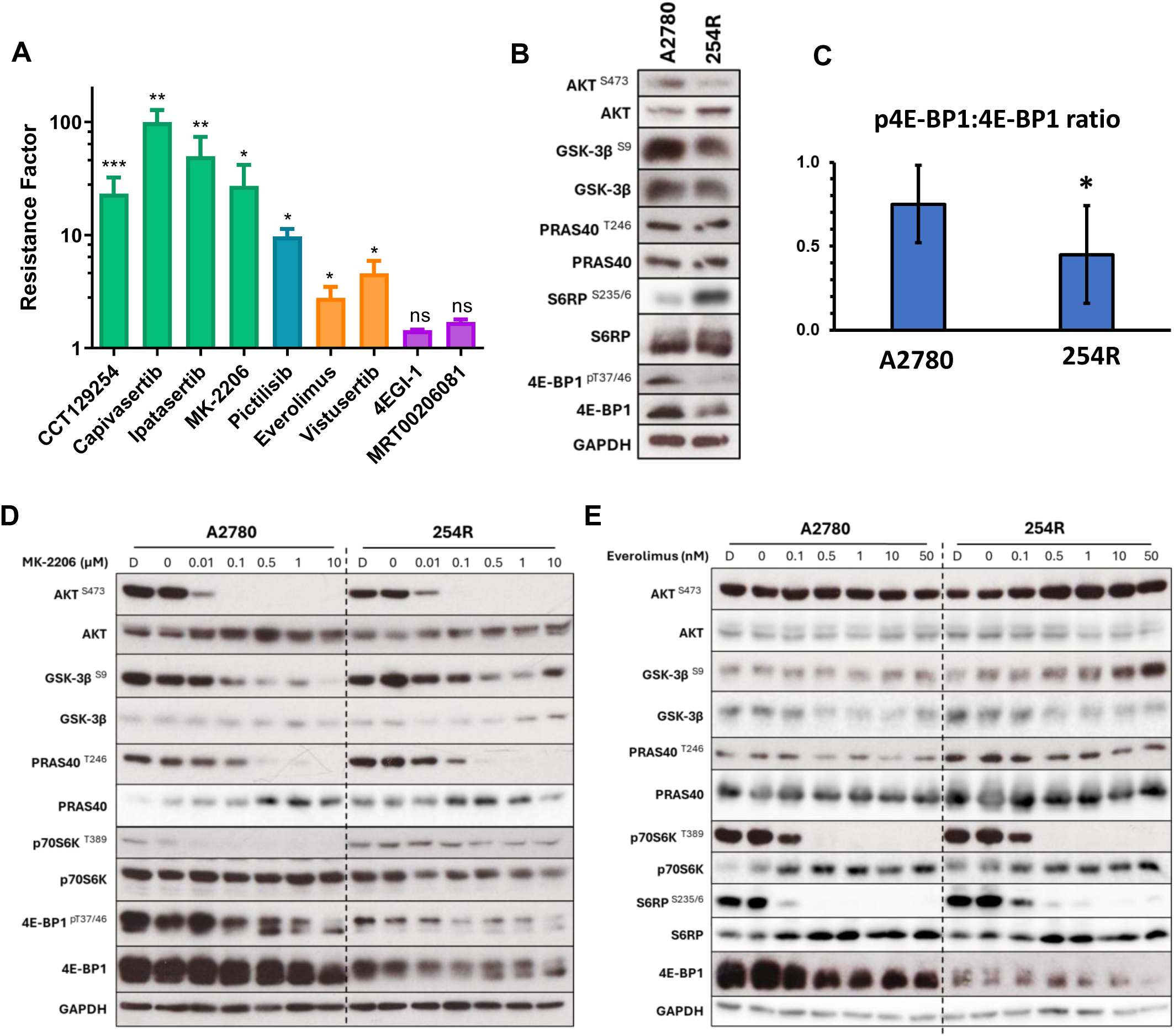
mTORC1 drives capivasertib resistance in a distinct role outside of the AKT/mTORC1 axis. **A)** Representation of the means ± standard deviations of resistance factors (RFs) from ≥3 independent experiments for the PAM pathway and protein synthesis inhibitors. Individual independent RF values were calculated as the ratio of 254R GI_50_ to A2780 GI_50_. Green; AKT, blue; PI3K, orange; mTORC and purple; protein synthesis inhibitors. **B)** Western blot of baseline signalling in the PAM pathway in A2780 and 254R cells with GAPDH as a loading control. **C)** Bar chart of the average ratio of 4E-BP1^T^^37/46^:total 4E-BP1. Band intensities were determined via ImageJ and normalized to the GAPDH loading control. **D)** PAM signalling in A2780 and 254R cells in response to increasing doses of MK-2206. The cells were grown for 72 hours with a final 24-hour treatment with MK-2206 prior to lysis and western blot analysis. E) PAM signalling in A2780 and 254R cells in response to increasing doses of everolimus. The cells were grown for 72 hours with a final 24-hour treatment with everolimus prior to lysis and western blot analysis. For western blots, representative western blots from ≥3 independent experiments are shown. The superscript indicates the phosphorylation site. D; DMSO control. Statistics: Unpaired t test with Welch’s correction *p<0.05, **p<0.01, ***p<0.001, ns; nonsignificant.

To determine whether resistance to AKT inhibitors in 254R cells results from reactivation of the PI3K/AKT/mTOR (PAM) signalling pathway, drug response assays were performed using the PI3K inhibitior pictilisib, the ATP-competitive mTORC1/2 inhibitor vistusertib, and the allosteric mTORC1 inhibitor everolimus. The 254R cells exhibited cross-resistance to each of these agents, with resistance factors of 9.8, 4.6, and 2.8, respectively (**Fig. 1A, Sup. Fig. 1A**). Given the observed resistance to mTORC1 inhibition, we next evaluated cap-dependent protein synthesis (CDPS) as a downstream vulnerability by treating cells with MRT00206081 and 4EGI-1, which are inhibitors of p70S6K and the eIF4E–4E-BP1 interaction, respectively. Notably, 254R cells did not display cross-resistance to either CDPS inhibitor (**Fig. 1A, Sup. Fig. 1A**), suggesting that targeting downstream effectors of mTOR may bypass the resistance mechanism.

Given the observed cross-resistance to mTORC1 inhibitors but not CDPS inhibitors, we hypothesised that the resistance mechanism in 254R cells may involve signalling downstream of mTORC1. To test this hypothesis, we examined key nodes of the PAM pathway proximal to and downstream of mTORC1 in A2780 versus 254R cells (**Fig. 1B**). Although a modest decrease in AKT phosphorylation at S473 was observed in 254R compared with A2780, the most notable changes occurred downstream of mTORC1. Specifically, 254R cells displayed a marked increase in the phosphorylation of the S6 ribosomal protein at S235/S236, which are phosphorylated by p70S6K, along with a reduction in both total 4E-BP1 expression and phosphorylation at T37/T46 (**Fig. 1B**), compared with A2780 cells. Since the ratio of phosphorylated 4E-BP1 to total 4E-BP1 influences the stoichiometry between 4E-BP1 and eIF4E, we quantified this value via densitometric analysis, which revealed a 1.67-fold reduction in the p-4E-BP1:total 4E-BP1 ratio in 254R relative to A2780 (**Fig. 1C**).

As both S6RP and 4E-BP1 are downstream of mTORC1, these baseline signalling alterations suggest elevated mTORC1 activity in 254R cells. To further investigate the dependency of mTORC1 signalling on AKT activity, we treated A2780 and 254R cells with increasing concentrations of the non-ATP competitive AKT inhibitor MK-2206 for 24 hours and evaluated PAM signalling (**Fig. 1D**). In both cell lines, MK-2206 induced a dose-dependent reduction in pS473-AKT, with complete loss of signal at 0.1 µM. Similarly, the phosphorylation of canonical AKT substrates, including pS9-GSK3β and pT246-PRAS40, was effectively suppressed in a concentration-dependent manner (**Fig. 1D**).

Despite efficient AKT pathway inhibition, the phosphorylation of p70S6K at T389, which is a direct target of mTORC1, remained resistant to MK-2206 in 254R cells, whereas it was abrogated at 0.01 µM in A2780 cells (**Fig. 1D**). Moreover, in addition to 4E-BP1 expression being reduced in 254R versus A2780, its phosphorylation at T37/T46 was more resistant to MK-2206, requiring higher concentrations to induce a shift from hyper to hypophosphorylated forms, in particular at 10 µM MK2206 (**Fig. 1D**). These data collectively indicate that in 254R cells, mTORC1 activity is uncoupled from upstream AKT signalling, indicating an AKT-independent mechanism of mTORC1 activation in acquired resistance.

To determine whether the MK-2206-resistant phosphorylation of mTORC1 downstream targets was still dependent on mTORC1 activity, A2780 and 254R cells were treated with increasing concentrations of the allosteric mTORC1 inhibitor everolimus for 24 hours, and PAM pathway signalling was assessed (**Fig. 1E**). Unlike MK-2206, everolimus did not reduce pS473-AKT levels. Instead, everolimus treatment led to a dose-dependent increase in pS473-AKT in both cell lines (**Fig. 1E**), a known feedback response to mTORC1 inhibition attributed to IRS-1 derepression ^33,34^.

Among all the PAM pathway markers examined, A2780 and 254R showed comparable responses to everolimus (**Fig. 1E**). Specifically, at 0.5 nM everolimus, the pT389-p70S6K signal was completely abolished, and a band shift in 4E-BP1 isoforms was observed in both cell lines, in sharp contrast to the partial effects observed with MK-2206 (**Fig. 1E**). These findings confirm that the phosphorylation of p70S6K and 4E-BP1 in 254R cells is dependent on mTORC1 activity and is not resistant to direct mTORC1 inhibition.

Together, these data suggest that capivasertib resistance in 254R cells may be driven by multiple resistance mechanisms. One such mechanism circumvents AKT function through hyperactivation of mTORC1 signalling, likely driven by a mechanism upstream or parallel to AKT but outside the canonical AKT–mTORC1 axis, as indicated by the inability of MK-2206 to inhibit pT389-p70S6K. Interestingly, despite showing clear MK-2206-resistant mTORC1 signalling, 254R cells retained their biochemical sensitivity to everolimus, although they exhibited statistically significant cross-resistance to its antiproliferative effects, as measured by GI_50_ determination (**Sup. Fig. 1A**). A potential second mechanism is through down-regulation of both total and phosphorylated 4E-BP1.

### Reduced phosphorylation and expression of 4E-BP1 in AKT inhibitor resistance

To resolve the conflicting observations regarding mTORC1 signalling in the 254R population, single-cell subclones were generated by limiting dilution. All eight subclones demonstrated significant resistance to both CCT129254 (6.3–11-fold) and capivasertib (16–54-fold), which was consistent with the parental 254R line (**Sup. Fig. 2A**). As with 254R, each subclone showed greater resistance to capivasertib than to CCT129254. Cross-resistance profiling revealed that all subclones were also resistant to the mTORC1 inhibitors everolimus (2.41–13.5-fold) and vistusertib (2.98–6.68-fold), with most showing greater resistance to vistusertib (**Sup. Fig. 2A**).Given our focus on acquired resistance to capivasertib, subclone 254R-B, which presented high capivasertib resistance combined with strong cross-resistance to other agents tested (**Sup. Fig. 2A**), was selected for further characterisation.

Consistent with the parental 254R population, 254R-B was resistant to all the AKT and mTORC1 inhibitors tested but remained sensitive to the CDPS inhibitor 4EGI-1 (**Fig. 2A**). Signalling analysis revealed minimal changes upstream of mTORC1 in 254R-B **(Fig. 2B)**. However, downstream signalling analysis revealed that phosphorylation of p70S6K at T389 increased in 254R-B versus A2780, independent of changes in total p70S6K, whilst both total and phosphorylated 4E-BP1 were reduced compared to A2780 (**Fig. 2B**). These altered baseline signalling features suggest converging mechanisms promoting resistant mTORC1 activity.

**Figure 2:**
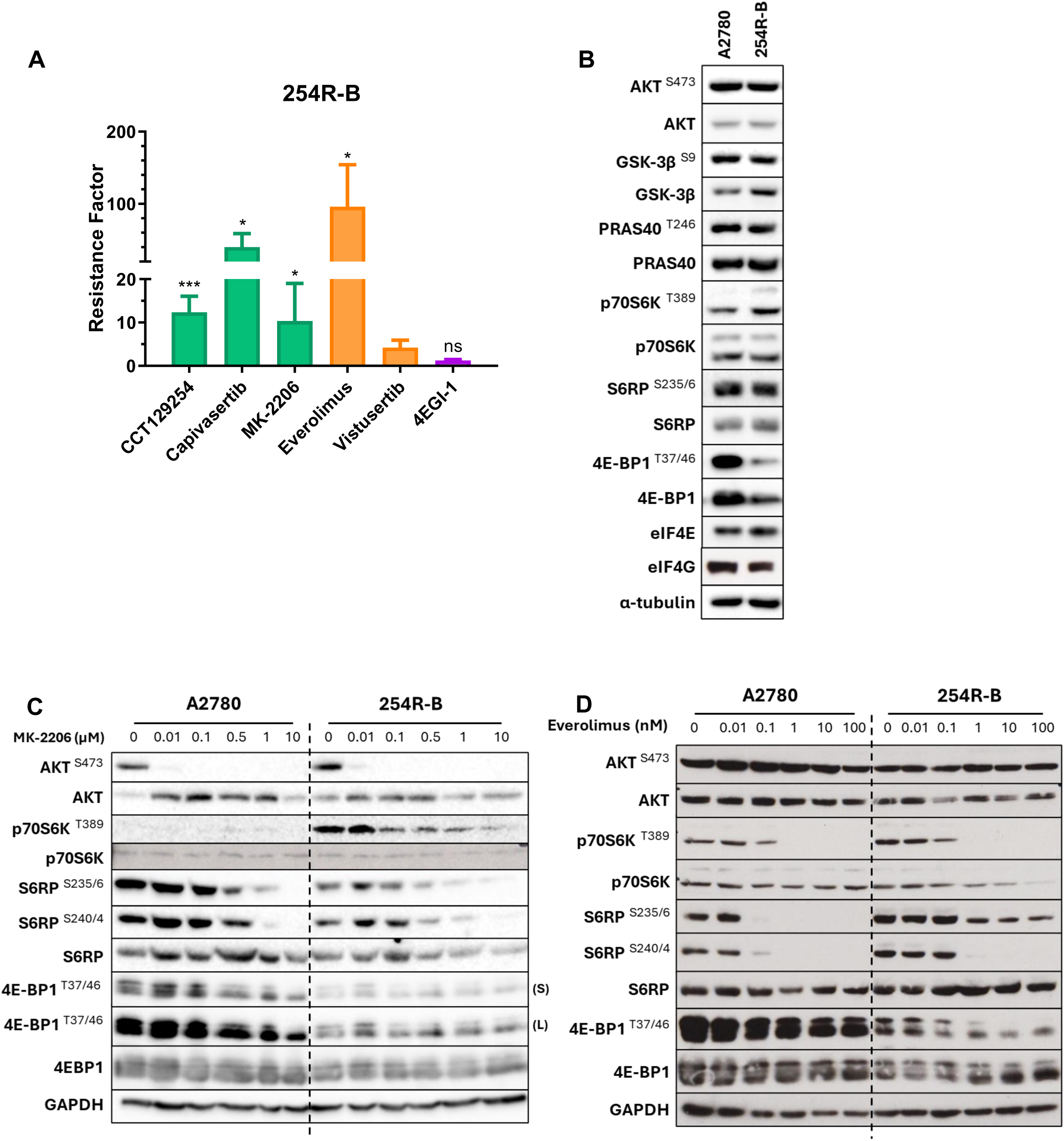
Cross-resistance profiling of subclone 254R-B to PAM inhibitors. **A)** Bar chart representing the mean ± standard deviation resistance factors for A2780 and 254R-B from ≥3 independent experiments. **B)** Western blot analysis of baseline signalling in the PAM pathway in A2780 and 254R-B cells, with α-tubulin as the loading control. **C)** Western blot of PAM signalling in A2780 and 254R-B cells in response to increasing doses of MK-2206 with GAPDH as a loading control. The cells were grown for 72 hours with a final 24-hour treatment with MK-2206 prior to lysis and western blot analysis. **D)** Western blot of PAM signalling in A2780 and 254R-B cells in response to increasing doses of everolimus, with GAPDH as a loading control. The cells were grown for 72 hours with a final 24-hour treatment with everolimus prior to lysis and western blot analysis. For western blots, representative western blots from ≥3 independent experiments are shown. The superscript indicates the phosphorylation site. Statistics: Unpaired t test with Welch’s correction *p<0.05, **p<0.01, ***p<0.001, ns; nonsignificant.

To assess mTORC1 signalling in response to AKT inhibition, A2780 and 254R-B cells were treated with increasing concentrations of MK-2206 for 24 hours (**Fig. 2C**). In both cell lines, pS473-AKT became undetectable at 0.1 μM MK-2206. While pT389-p70S6K was undetectable in A2780 cells, it appeared to be again elevated at baseline in 254R-B and decreased in a dose-dependent manner in the presence of MK-2206. Both A2780 and 254R demonstrated a dose-dependent decrease in S235/236 and S240/244 signals in the presence of MK-2206 (**Fig. 2C**). Additionally, phosphorylated 4E-BP1 was lower in 254R-B-treated cells than in A2780-treated cells at baseline, and both showed a dose-dependent decrease in phosphorylation in the presence of MK-2206. Collectively, these findings indicate that despite increased baseline p70S6K activity, mTORC1 signalling in 254R-B remains sensitive to AKT inhibition.

Given the high degree of everolimus resistance observed in 254R-B, we next examined the effect of this drug on downstream AKT/mTORC1 signalling (**Fig. 2D**). In both A2780 and 254R-B cells, everolimus treatment led to a dose-dependent reduction in pT389-p70S6K and S6RP phosphorylation at S235/236 and S240/244, with near-complete suppression of these signals by 1 nM. However, S6RP pS235/236 remained detectable in 254R-B even at the highest dose (100 nM), with a clear reduction between 0.1 and 1 nM. The baseline pT37/46-4E-BP1 level was reduced in 254R-B, and both cell lines shifted from high-molecular-weight (hyperphosphorylated) to low-molecular-weight (hypophosphorylated) 4E-BP1 isoforms with increasing everolimus concentrations. Notably, pT37/46 4E-BP1 phosphorylation persisted in the high-molecular-weight isoforms at 1 nM everolimus, unlike pT389-p70S6K, which is consistent with prior observations that rapalogues more effectively inhibit mTORC1 regulation of p70S6K than 4E-BP1^35^. Importantly, neither p70S6K nor 4E-BP1 phosphorylation was more resistant to everolimus in 254R-B than in A2780. In contrast, S6RP phosphorylation at S235/236 in 254R-B did show resistance to inhibition with everolimus, indicating that part of this signal may be maintained independently of mTORC1.

In summary, 254R-B displays altered mTORC1 signalling dynamics in response to PAM pathway inhibitors. While mTORC1 activity remains sensitive to MK-2206, 4E-BP1 phosphorylation and expression are reduced and less functionally responsive, regardless of AKT inhibition. Furthermore, 254R-B exhibited persistent S6RP S235/236 phosphorylation despite everolimus treatment, a feature not observed with AKT inhibition, suggesting that distinct mTORC1 downstream outputs may be differentially regulated in the context of acquired resistance.

### Reduced 4E-BP1-eIF4E interaction in 245R-B

Clone 254R-B exhibited resistance to AKT and mTORC1 inhibitors but remained sensitive to the eIF4F complex inhibitor 4EGI-1 (**Fig. 2A**). This finding, combined with reduced 4E-BP1 expression and increased S6RP phosphorylation compared with those in the A2780 cell line, suggested that the resistance phenotype in 254R-B cells may converge at the level of cap-dependent protein synthesis (CDPS).

The canonical function of 4E-BP1 is to bind and sequester eIF4E, the mRNA cap-binding protein required for CDPS (**Fig. 3A**). To assess whether this interaction was altered in 254R-B, we performed a 7-methyl-GTP (m7GTP) pull-down assay following 24 hours of treatment with PAM inhibitors targeting mTORC1 directly or indirectly. Since only hypophosphorylated 4E-BP1 can bind eIF4E, a single 4E-BP1 band was observed in the pull-down data (**Fig. 3B**).

**Figure 3:**
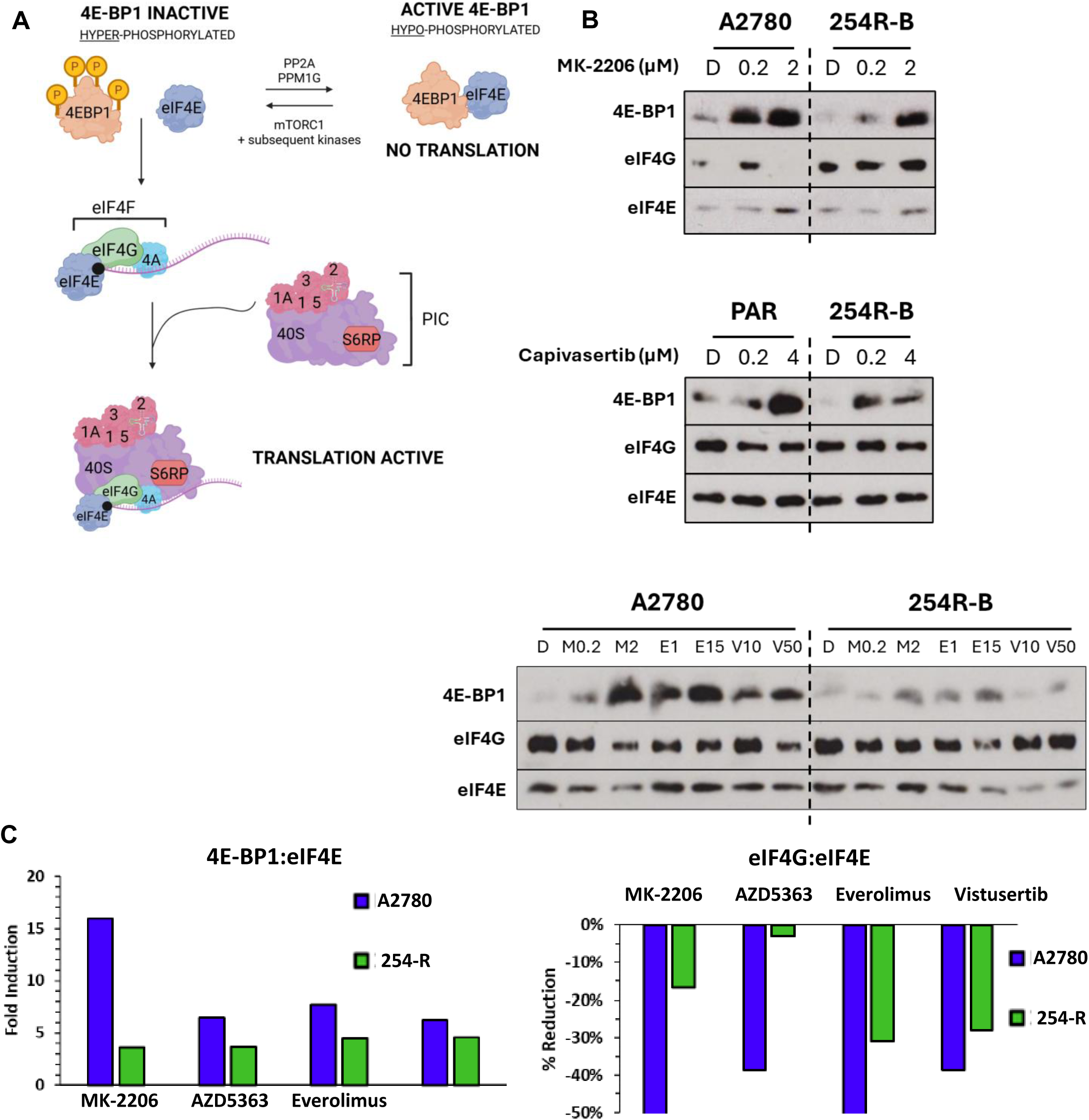
Reduced 4E-BP1/eIF4E interaction in 245R-B but no impact on global protein synthesis or increased cap-dependent protein synthesis. **A)** Schematic of the role of 4E-BP1 and S6RP in cap-dependent protein synthesis. When 4E-BP1 is hypophosphorylated, it binds and sequesters the mRNA cap-binding protein eIF4E . 4E-BP1 becomes inactivated through phosphorylation at multiple residues and cannot interact with eIF4E. eIF4E brings the mRNA to the scaffold protein eIF4G (4G) and, with eIF4A (4A), forms the eIF4F initiation complex. This allows subsequent recruitment of the preinitiation complex (PIC) for ribosomal docking. S6RP is bound to the 40S ribosomal subunit. **B)** Dynamics of 4E-BP1 and eIF4G binding to eIF4E with PAM inhibitor treatment in A2780 and 254R-B cells. Cells treated with either MK-2206 (top), capivasertib (middle), MK-2206 (M; µM), everolimus (E; nM), or vistusertib (V; nM) (lower) at A2780 GI_50_ for 24 hours were subjected to a 7-methyl-GTP (m7GTP) pull-down assay followed by western blot analysis for the proteins indicated. **C)** Quantitative analysis of the dynamics of 4E-BP1 and eIF4G binding to eIF4E with PAM inhibitor treatment in A2780 and 254R-B cells. Western blots from **(B)** were used to quantify the band intensities determined with ImageJ and adjusted to eIF4E loading. Induction was calculated as the fold difference between the adjusted area band intensity of 4E-BP1 in the DMSO control versus the maximum drug concentration per drug. The percentage reduction in the eIF4G:eIF4E ratio was calculated as the percentage difference between the adjusted area band intensity of eIF4G in the DMSO control versus the maximum drug concentration per drug.

Under AKT inhibition with MK-2206, 254R-B exhibited lower baseline 4E-BP1–eIF4E binding than A2780 did. Treatment with MK-2206 increased association between 4E-BP1 and eIF4E in both cell lines; however, the induction was more robust in A2780 (16-fold) than in 254R-B (4-fold) (**Fig. 3B, 3C**). Correspondingly, eIF4G association with eIF4E was significantly reduced in A2780 cells (57% reduction), whereas 254R-B showed only a 17% decrease. Capivasertib treatment yielded similar trends. In A2780 cells, association between 4E-BP1 and eIF4E increased 6.5-fold, but was reduced for eIF4G by 39%. In contrast, 254R-B resulted in only a 3.7-fold increase in association between 4E-BP1 and eIF4E and a minimal (2.9%) reduction in eIF4G association (**Fig. 3B, 3C**).

Similar results were obtained with the mTORC1 inhibitors everolimus and vistusertib. In A2780 cells, both drugs increased 4E-BP1 and eIF4E association (7.7-fold and 6.2-fold, respectively) and reduced association of eIF4G and eIF4E by 66.3% and 39%, respectively. However, in 254R-B, 4E-BP1 association with eIF4E increased to a lesser extent (∼4.5-fold for both drugs), and eIF4G association with eIF4E was reduced by only 30% at the highest drug concentrations (**Fig. 3B, 3C**). Together, these results indicate that, compared with A2780, 4E-BP1 in 254R-B is less responsive to phosphorylation inhibition, resulting in impaired displacement of eIF4G from the eIF4F complex.

The observed changes in the ratio of 4E-BP1 or eIF4G binding with eIF4E may suggest that CDPS is altered in 254R-B. To evaluate whether altered 4E-BP1/eIF4G dynamics affect CDPS, we used a bicistronic dual-luciferase reporter assay (pRL-IRES-FL), in which Renilla luciferase reports CDPS and firefly luciferase reflects cap-independent translation (CIPS) via the poliovirus IRES element (**Fig. 4A**). Compared with A2780, 254R-B resulted in a striking 2400% increase in Renilla activity and an 83% decrease in firefly activity (**Fig. 4B**), corresponding to a 9-fold increase in the CDPS:CIPS ratio and suggesting enhanced CDPS (**Fig. 4C**).

**Figure 4:**
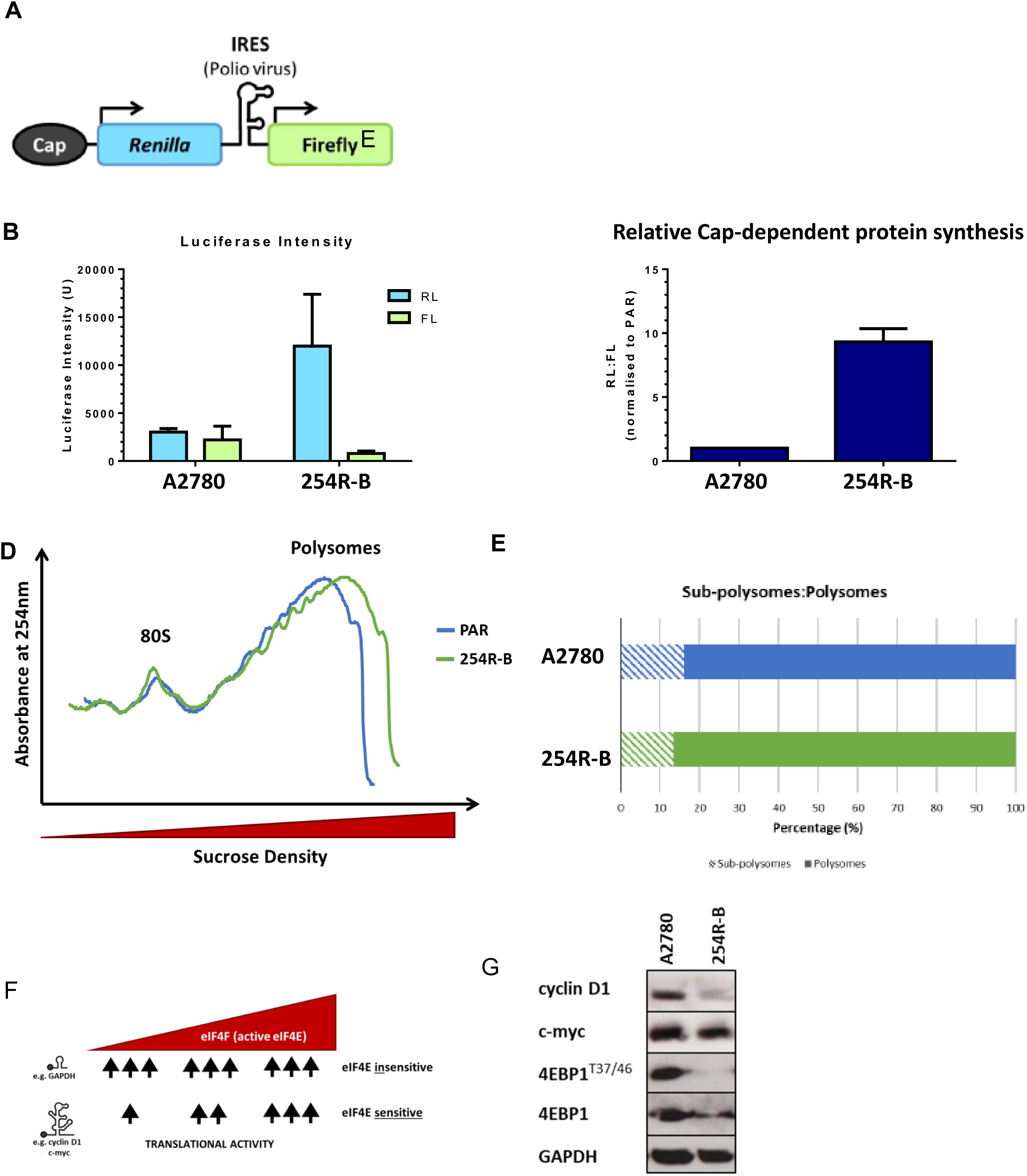
Induction of cap-dependent protein synthesis occurs in 254R-B versus A2780 cells. **A)** Schematic of the pRL-IRES-FL plasmid. **B)** Determination of the relative levels of cap-dependent and IRES-dependent protein synthesis in A2780 and 254R-B cells. **C)** Relative cap-dependent protein synthesis in A2780 versus 254R-B cells was calculated as the ratio of the luciferase intensity of Renilla luciferase (RL) to that of firefly luciferase (FL). **D)** Investigation of global protein synthesis in the A2780 and 254R-B cell lines. The absorbance of sucrose gradient fractions was analysed at 254 nm to produce a trace, as shown, and labelled as indicated (a representative example of three independent experiments). Increased absorbance is correlated with increased levels of the relevant mRNAs in the analysed fraction. (**E**) Bar graph of average polysome and subpolysome percentages. The areas under the polysome and subpolysome curves were calculated and analysed as a percentage of the total area under the subpolysome and polysome peaks (average of three independent experiments). **F)** Schematic of the protein synthesis activity of eIF4E-sensitive mRNAs (top row), which are strongly influenced by changes in eIF4E availability, whereas eIF4E-insensitive mRNAs (bottom row), in contrast, are marginally influenced by eIF4E availability. Illustrations based on and adapted from Roux and Topisirovic (2012) and Leppek et al. (2018)^40,56^. **G)** Western blot analysis of the baseline expression of cyclin D1, c-myc, 4E-BP1 and 4E-BP1^T^^37/46^ in the A2780 and 254R-B-cell lines, with GAPDH as a loading control (representative of ≥3 independent experiments). The superscript indicates the phosphorylation site.

We next asked whether this increase in CDPS was associated with global increases in protein synthesis. Polysome profiling was performed via the use of cycloheximide to freeze elongating ribosomes. Actively translated mRNAs (polysomes) were detected in heavy fractions, whereas free ribosomes and poorly translated mRNAs (subpolysomes) were detected in lighter fractions (**Fig. 4D**). Quantification of the area under the curve revealed a modest decrease in the proportion of subpolysomes in 254R-B (13.4%) compared with A2780 (16.1%) and a corresponding slight increase in polysomes (86.6% vs. 83.9%), suggesting a limited increase in global translation (**Fig. 4E**).

These findings suggest that elevated CDPS in 254R-B may not reflect a global increase in protein synthesis but rather a selective increase in the translation of eIF4E-sensitive transcripts. These transcripts, including cyclin D1 and c-Myc, possess long, structured 5′ UTRs that require the eIF4F complex, particularly the helicase activity of eIF4A, for efficient translation (**Fig. 4F**) ^36–40^. However, despite increased CDPS, the protein expression of cyclin D1 was lower in 254R-B versus A2780, whilst c-myc was similar in both cell lines (**Fig. 4F**), suggesting that these transcripts are not the primary drivers of the CDPS phenotype and may be suppressed by other regulatory mechanisms (**Fig. 4G**) ^36–40^.

### Ectopic expression of wild-type or constitutively active 4E-BP1 restores capivasertib sensitivity

To evaluate whether restoration of 4E-BP1 activity could sensitise 254R-B cells to capivasertib, we overexpressed both wild-type (WT) and constitutively active mutant forms of 4E-BP1. The 254R-B cells were transfected with pcDNA3-4E-BP1WT-HISMYC (encoding wild-type human 4E-BP1) or pcDNA3-4E-BP15A-HISMYC (encoding the nonphosphorylatable, constitutively active 5A mutant) ^41,42^. The 5A mutant harbours alanine substitutions at five phosphorylation sites (T36, T47, T70, S83, and S65), preventing its inactivation by phosphorylation and rendering it constitutively active.

Transient expression was confirmed via His-tag detection (**Fig. 5A**). In WT-transfected cells, both total and phosphorylated 4E-BP1 (T37/46) levels increased, whereas 5A-transfected cells presented increased total 4E-BP1 without detectable T37/46 phosphorylation, as expected (**Fig. 5A**). Stable 254R-B sublines expressing WT or 5A 4E-BP1 were generated via the use of G418 resistance cassettes. Following antibiotic selection, both the clonal and polyclonal populations were expanded and validated via western blotting (**Sup. Fig. 3**). The His-Myc tag caused exogenous 4E-BP1 to migrate more slowly, allowing clear separation from the endogenous protein. All clones except WT 7 and 1P successfully expressed the transgene. WT clones 14 and 17 and 5A clone 4 were selected for further analysis. Notably, compared with the untransfected 254R-B cell line, clone 4 (here after referred to as 5A4) presented reduced endogenous 4E-BP1 phosphorylation.

**Figure 5:**
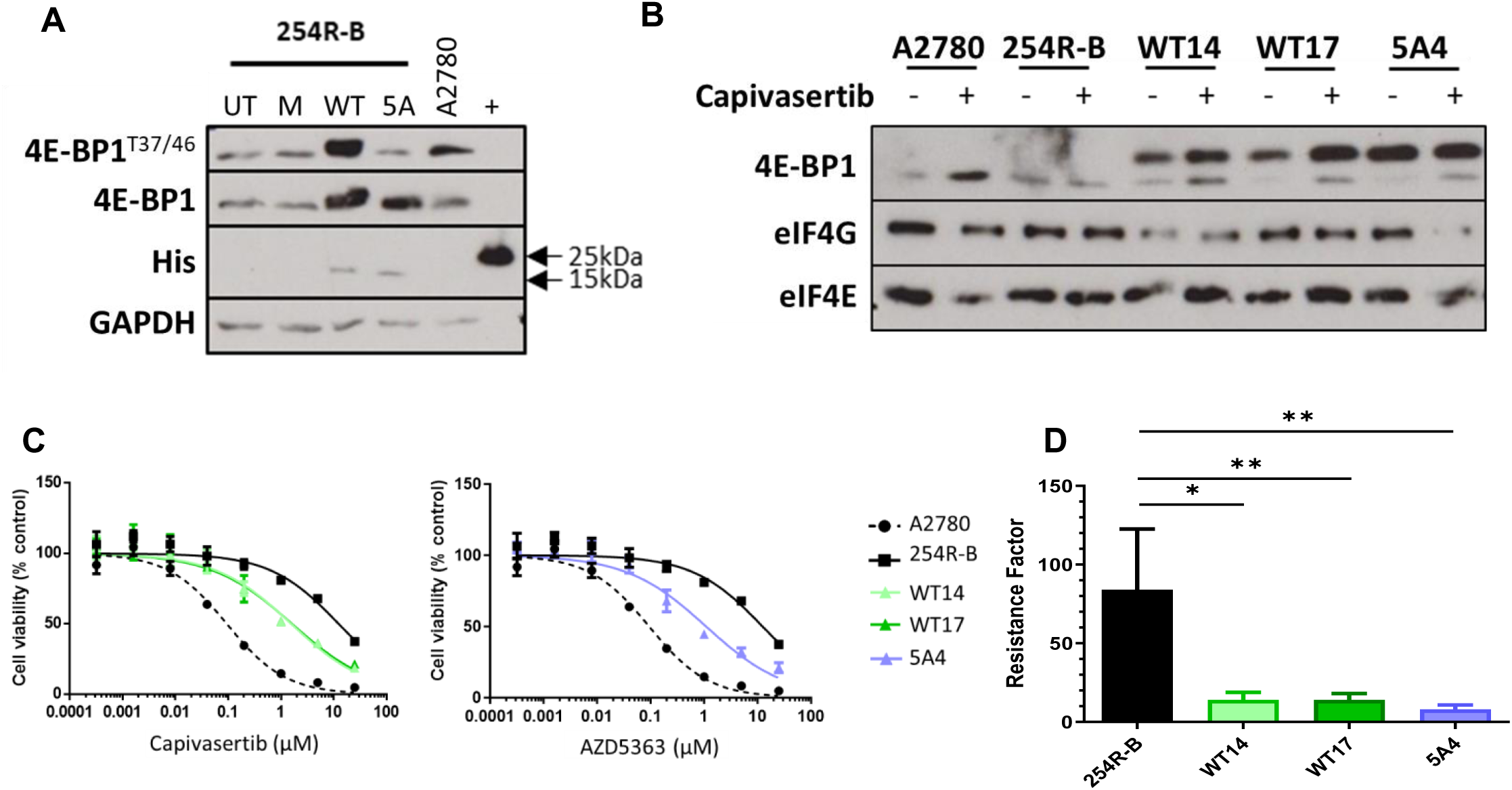
Expression of exogenous wild-type and constitutively active 4E-BP1 reduces capivasertib resistance. **A)** Transient expression of wild-type and 5A-mutant 4E-BP1 in 254R-B cells. Forty-eight hours post-transfection, the cells were lysed and analysed by western blotting for the proteins indicated, with GAPDH as a loading control. A2780: untransfected; His: His-tagged protein used as a positive control. **B)** The indicated cell lines were treated with capivasertib (4 μM) for 24 hours, and lysates were prepared and subjected to a 7-methyl-GTP (m7GTP) pull-down assay followed by western blot analysis for the proteins indicated. Ex: exogenous 4E-BP1; En: endogenous 4E-BP-1. **C)** Representative dosecresponse curves for capivasertib in the indicated cell lines. After 48 hours, the cells were plated and treated with serially diluted concentrations of capivasertib, followed by incubation for 96 hours, and then, an SRB assay was performed, and the cell viability was calculated versus that of untreated cells for each cell line from ≥3 independent experiments. **D)** Bar chart representing the mean ± standard deviation resistance factors for capivasertib in the indicated cell lines from ≥3 independent experiments. **E)** Capivasertib resistance factor (from **D**) against the relative density of 4E-BP1 expression with capivasertib treatment (+) (from **B**). Band intensities were determined via ImageJ and adjusted relative to eIF4E. The ratio of p4E-BP1:4E-BP1 for each cell line within brackets was calculated via densitometric analysis as described above (relative to GAPDH) for the bands in supplementary figure 3. Statistics: Unpaired t test with Welch’s correction *p<0.05, **p<0.01, ***p<0.001, NS; nonsignificant.

To assess the functional activity of 4E-BP1 in these sublines, m^7^GTP pull-down assays were performed to examine interactions with eIF4E and eIF4G (**Fig. 5B**). In A2780 cells, capivasertib treatment increased 4E-BP1 association with eIF4E and decreased eIF4G association. In contrast, 254R-B cells presented no change in 4E-BP1 or eIF4G association with eIF4E following drug treatment.

Compared with A2780 or 254R-B, all 254R-B 4E-BP1-overexpressing sublines presented high basal levels of exogenous 4E-BP1 association with eIF4E and lower association between eIF4G and eIF4E, indicating enhanced 4E-BP1 functional activity (**Fig. 5B**). When treated with capivasertib, WT14 and WT17 presented increased association of exogenous 4E-BP1 with eIF4E versus untransfected 254R-B. This effect was not observed for the 5A4 subline expressing the non-phosphorylatable, constitutively active form of 4E-BP1, as expected due to the mutated form (**Fig. 5B**). However, 5A4 still presented reduced eIF4G–eIF4E binding upon capivasertib treatment (**Fig. 5B)**. These findings suggest that exogenous 4E-BP1 expression partially restored the sensitivity of 4E-BP1–eIF4E binding to capivasertib treatment in 254R-B cells.

Functionally, overexpression of either WT or 5A4 4E-BP1 significantly sensitised 254R-B cells to capivasertib. WT14, WT17, and 5A4 cells presented markedly lower resistance factors (7.6–14-fold) than the 254R-B cell line (84.0-fold) (**Figs. 5C, 5D**).

Taken together, these results demonstrate that exogenous expression of either wild-type or constitutively active 4E-BP1 enhances 4E-BP1 functional activity and restores capivasertib sensitivity in 254R-B cells, highlighting a role for impaired 4E-BP1 regulation in the resistance phenotype.

## Discussion

The PI3K/AKT/mTOR (PAM) pathway is among the most frequently dysregulated signalling cascades in human cancers, where its hyperactivation drives proliferation, survival, and therapy resistance ^1^. Consequently, it has become a central focus of molecularly targeted therapy development, with both PI3 kinase and AKT inhibitors, such as apelisib and capivasertib, respectively, now approved for cancer treatment ^15,43^. Despite these advances, the clinical benefit of PAM pathway inhibition is often transient, as many patients relapse due to acquired resistance^20,21,44^ Understanding the molecular mechanisms of resistance is critical for informing rational combination strategies and for developing predictive biomarkers to guide therapy.

To investigate the mechanisms of resistance to AKT-targeted therapies, we generated an AKT inhibitor-resistant ovarian cancer cell line, A2780 254R, via chronic exposure to the early capivasertib analogue CCT129254. This model exhibited robust resistance not only to CCT129254 but also to capivasertib itself, validating its utility as a tool for dissecting resistance mechanisms relevant to clinical inhibitors (**Fig. 1A, Sup Fig. 1A**). Mechanistic analysis revealed that resistance to 254R was not attributable to activating mutations in AKT isoforms or increased AKT kinase activity (**Sup Fig. 1B, C)**. Instead, resistance was associated with increased mTORC1 signalling, as evidenced by the sustained phosphorylation of canonical mTORC1 substrates despite AKT inhibition (**Fig. 1D**).

Interestingly, 254R cells also displayed cross-resistance to PI3K and mTOR inhibitors, suggesting broad rewiring of the PAM axis. However, sensitivity to cap-dependent protein synthesis (CDPS) inhibitors remains largely intact. Notably, although 254R cells exhibited increased mTORC1 activity, they remained somewhat sensitive to the mTORC1 inhibitor everolimus (**Fig. 1E**), prompting further investigation into the apparent disconnect between mTORC1 activation and the drug response. To explore the possibility of intratumoral heterogeneity in resistance mechanisms, we derived eight single-cell subclones from the 254R population.

All eight subclones maintained resistance to CCT129254, although they exhibited variable sensitivity to capivasertib and everolimus. Among the sub-clones, 254R-B displayed pronounced resistance to both capivasertib and everolimus, prompting its selection for detailed mechanistic analysis.In 254R-B, baseline phosphorylation of p70S6K (T389), a key mTORC1 substrate, was greater than that in the A2780 parental cell line and decreased following AKT inhibition, indicating persistent mTORC1 signalling with sensitivity to AKT inhibition (**Fig. 2C**). However, baseline downstream S6 ribosomal protein (S235/236, S240/244) phosphorylation in 254R-B did not mirror this increase and indeed was reduced compared with that in A2780, suggesting altered substrate specificity of p70S6K or compensation by other kinases. Treatment with everolimus effectively suppressed p70S6K phosphorylation but failed to fully abolish S6RP phosphorylation beyond 1 nM, implicating additional kinases, such as p90RSK, in maintaining residual S6RP activity (**Fig. 2D**). This everolimus-insensitive S6RP phosphorylation further supports the notion that noncanonical signalling contributes to resistance. In contrast, the allosteric AKT inhibitor MK-2206 suppressed both p70S6K and S6RP phosphorylation (**Fig. 2C**), suggesting differential pathway dependencies across inhibitor classes.

To assess whether dysregulated cap-dependent translation contributes to resistance, we performed mcGTP cap-binding assays to evaluate complex formation between eIF4E, 4E-BP1, and eIF4G. While AKT and mTORC1 inhibition enhanced the 4E-BP1–eIF4E interaction in both parental and resistant cells, this interaction was markedly reduced in 254R-B cells, which was consistent with decreased 4E-BP1 expression and phosphorylation. Moreover, drug-induced disruption of eIF4G binding to eIF4E was minimal in resistant cells, indicating the persistence of active eIF4F complexes (**Fig. 3B, C**). A dual-luciferase reporter assay confirmed an ∼9-fold increase in basal CDPS activity in 254R-B cells relative to parental cells (**Fig. 4B, C**), suggesting a translationally permissive state despite global protein synthesis and the expression of the canonical eIF4E target c-myc remaining unchanged or, in the case of cyclin D1, being dramatically reduced in 254R-B cells. This selective translational rewiring highlights a nuanced layer of resistance regulation.

Crucially, forced expression of either wild-type 4E-BP1 or a constitutively active, nonphosphorylatable mutant (5A4) restored capivasertib sensitivity in 25R-B cells versus A2780 parental cells, reducing the resistance factor by more than 10-fold (**Fig. 5C-E**). This functional rescue directly implicates impaired 4E-BP1 activity in driving the resistant phenotype.

Our findings align with results from a recent genome-wide CRISPR screen in PTEN-deficient breast cancer cells, which identified TSC1, TSC2, and STK11, all key negative regulators of mTORC1 and AMPK, as top resistance hits to capivasertib ^21^. Additional hits included GATOR1 complex members (DEPDC5, NPRL2, DDIT4), further implicating deregulated nutrient sensing and mTORC1 hyperactivation as mechanisms of resistance to capivasertib. Similarly, loss of LCMT1, a regulator of PP2A phosphatase activity, has been shown to sustain 4E-BP1 and p70S6K phosphorylation in the context of AKT inhibition, echoing our findings in 254R-B ^45^.

Together, these results support a model in which resistance to AKT inhibition converges on deregulated mTORC1 signalling and impairs 4E-BP1 function, leading to enhanced cap-dependent translation (**Figure 6**). This rewired translational landscape may allow tumour cells to sustain the expression of prosurvival and growth-related proteins despite upstream pathway inhibition. This finding suggests that therapeutic resistance may emerge through adaptive increases in eIF4F activity, even in the absence of overt changes in mTORC1 output.

**Figure 6:**
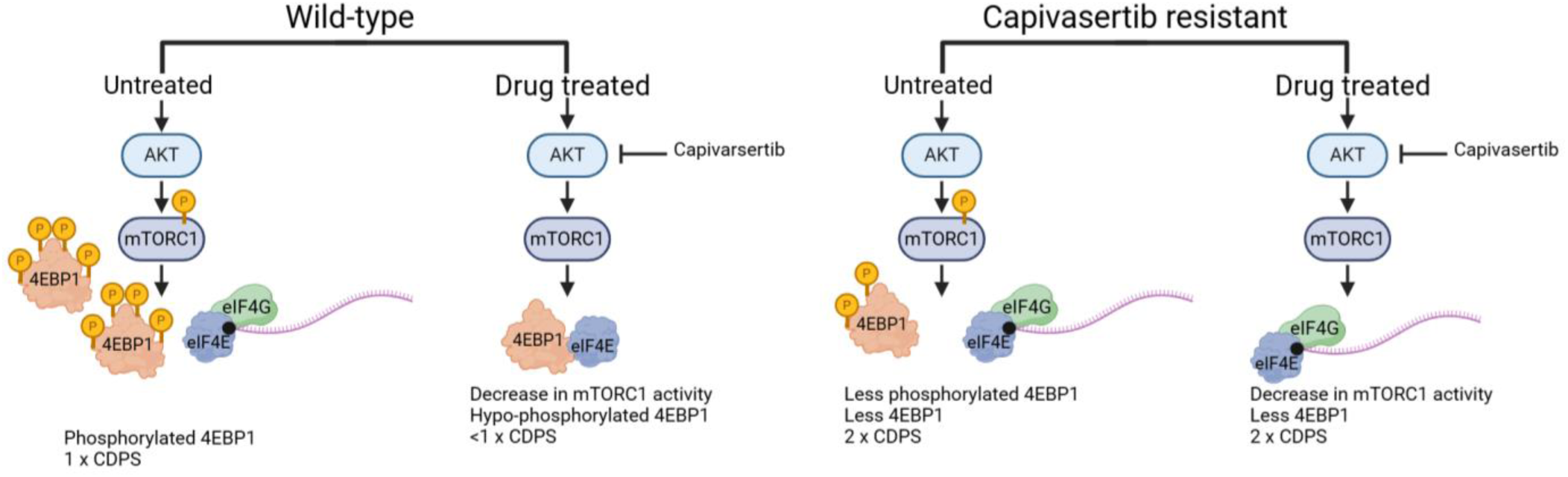

Importantly, eIF4E overexpression and hyperphosphorylation have been reported across several tumour types, including NSCLC, prostate cancer, melanoma, and ovarian cancer, highlighting the broader relevance of translational control in cancer progression and drug resistance ^46^. Thus, monitoring the expression and phosphorylation of 4E-BP1 and eIF4E may provide predictive biomarkers of resistance to AKT-targeted therapies.

Despite extensive interest, targeting the eIF4F complex in the clinic remains challenging. The eIF4E antisense oligonucleotide LY2275796 demonstrated target knockdown in preclinical models but yielded limited clinical benefit because of incomplete inhibition of translation ^47^. Ribavirin, which disrupts eIF4E-cap interactions, shows initial activity in acute myeloid leukaemia but has poor pharmacokinetics and limited tumour uptake ^48–50^. More recently, compounds such as 4Ei-1 and 4EGI-1, which both inhibit the eIF4E–eIF4G interaction, have demonstrated promising activity in preclinical models of lung, breast, and multiple myeloma, but their clinical development is still underway ^51–55^.

In summary, our study has revealed that reduced 4E-BP1 function and elevated cap-dependent protein synthesis are central drivers of acquired resistance to capivasertib in ovarian cancer cells. These findings underscore the translational machinery as a critical node of resistance and therapeutic vulnerability downstream of mTORC1. In the future, pharmacological targeting of the eIF4F complex may represent a promising strategy to overcome resistance and enhance the durability of AKT-targeted therapies. Biomarker-driven patient selection on the basis of 4E-BP1/eIF4E axis activity could further refine therapeutic approaches in this context.

## Acknowledgements

We would like to thank Dr. Simon Cook (Babraham Institute, Cambridge) for the kind gift of the bicistronic pRL-IRES-FL plasmid; Dr. Chris Proud (SAHMRI, Australia) for the kind gifts of pcDNA3-4E-BP1WT-HISMYC (human wild-type 4E-BP1; WT) and pcDNA3-4E-BP15A-HISMYC (human 5A-mutant 4E-BP1; 5A); Andy Merritt and Ed McIver for MRT00206081; and John Caldwell and Ian Collins (Institute of Cancer Research) for the kind gift of CCT129254. Figures were created by BioRender.com.

## Sources of Support/Funding

HEG, JMJ, DTA, and MDG all acknowledge funding from the University of Kent. EB and MDG acknowledge funding from the EB Hutchinson Charitable Trust. DTA and MDG also acknowledge funding from The Institute of Cancer Research (ICR) and in particular the MRC Doctoral Training Program at the ICR. Additional funding was received from Kent Cancer Trust (HEG).

---

**Supplementary Figure 1:**
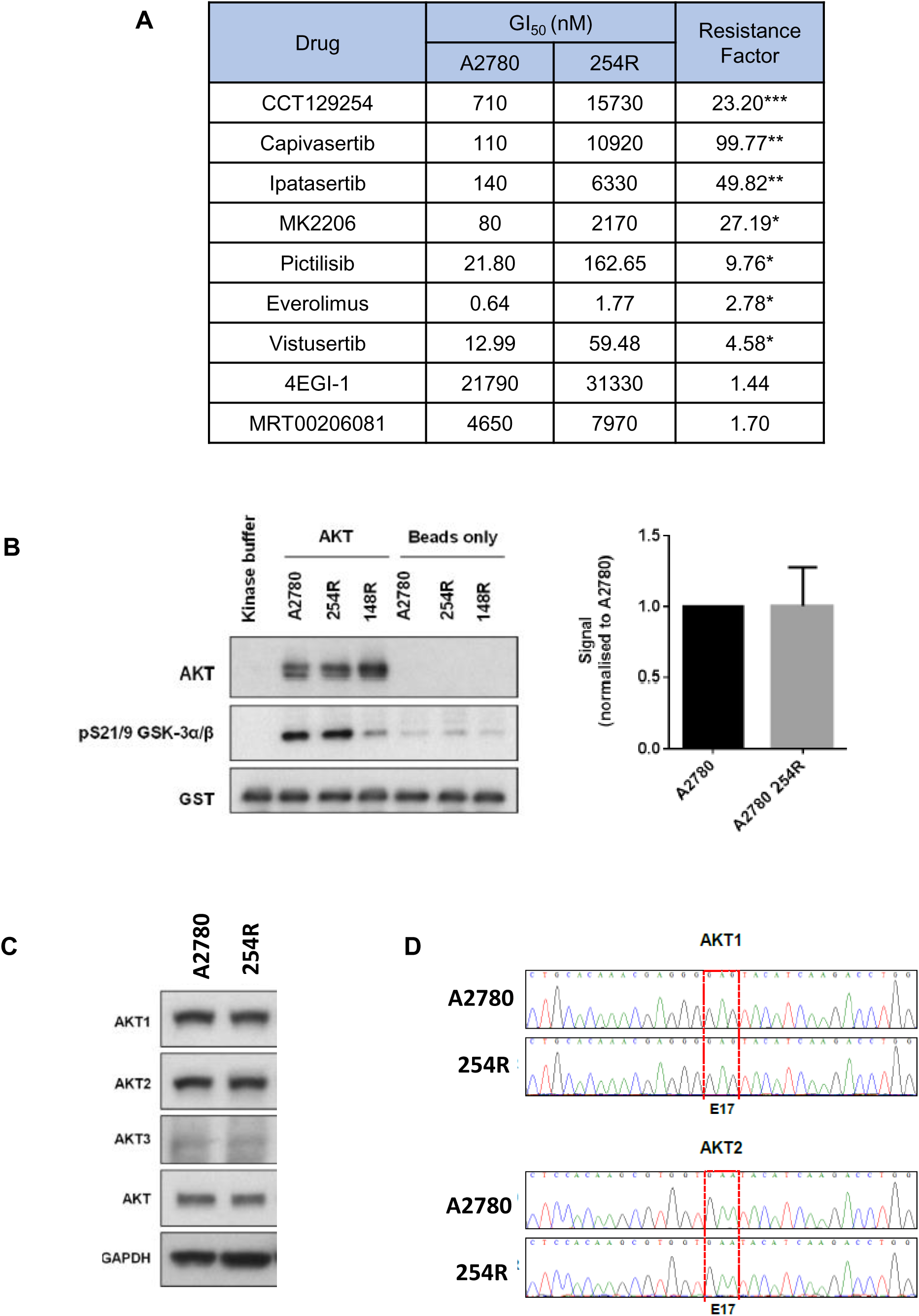
Increases in AKT expression and mutations are not responsible for the resistance mechanism of 254R. **A)** Summary of GI_50_ values and resistance factors (RFs) in A2780 and 254R cells. GI_50_ values represent the means from ≥3 independent experiments. Individual independent RF values were calculated as the ratio of 254R GI_50_ to A2780 GI_50_. **B)** Determination of AKT activity by in vitro kinase assay. Left: representative western blot analysis of the *in vitro* kinase assay. The antibody against pS21/9 GSK-3α/β detects the phosphorylation of the substrate fusion protein. Kinase buffer lane contained the kinase reaction mixture without addition of immunoprecipiates. 148R is another cell line not relevant to this paper. Right: Densiometric analysis of western blots **C)** Western blot analysis of AKT isoform expression. The data are representative of two independent experiments. **D)** Sequence traces of AKT isoforms. Dideoxynucleotide sequencing was performed on AKT1 and AKT2 reverse transcribed mRNAs extracted from A2780 and A2780 254R. Traces depict codons for glutamic acid 17 (highlighted in the red box) and surrounding nucleotides. The sequences were analysed via the BioEdit Sequence Alignment Editor v 7.1.3.0.

**Supplementary Figure 2:**
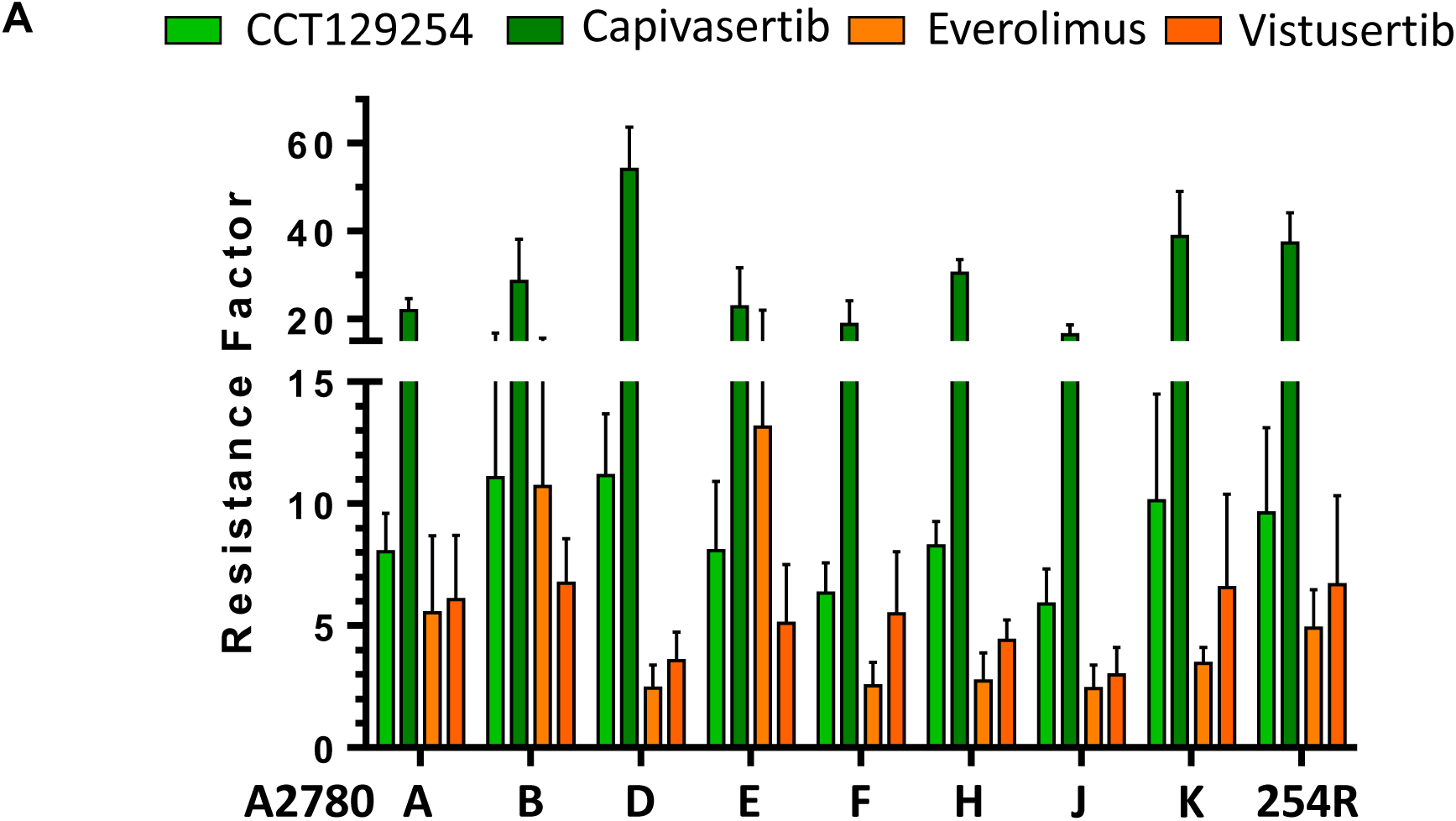
Screening of A2780 254R clones for cross-resistance and baseline mTORC1 signalling. **A)** Bar chart summarizing the means ± standard deviations of resistance factors for A2780 and 254R subclones; green, AKT; orange, mTORC inhibitors from ≥3 independent experiments.

**Supplementary Figure 3:**
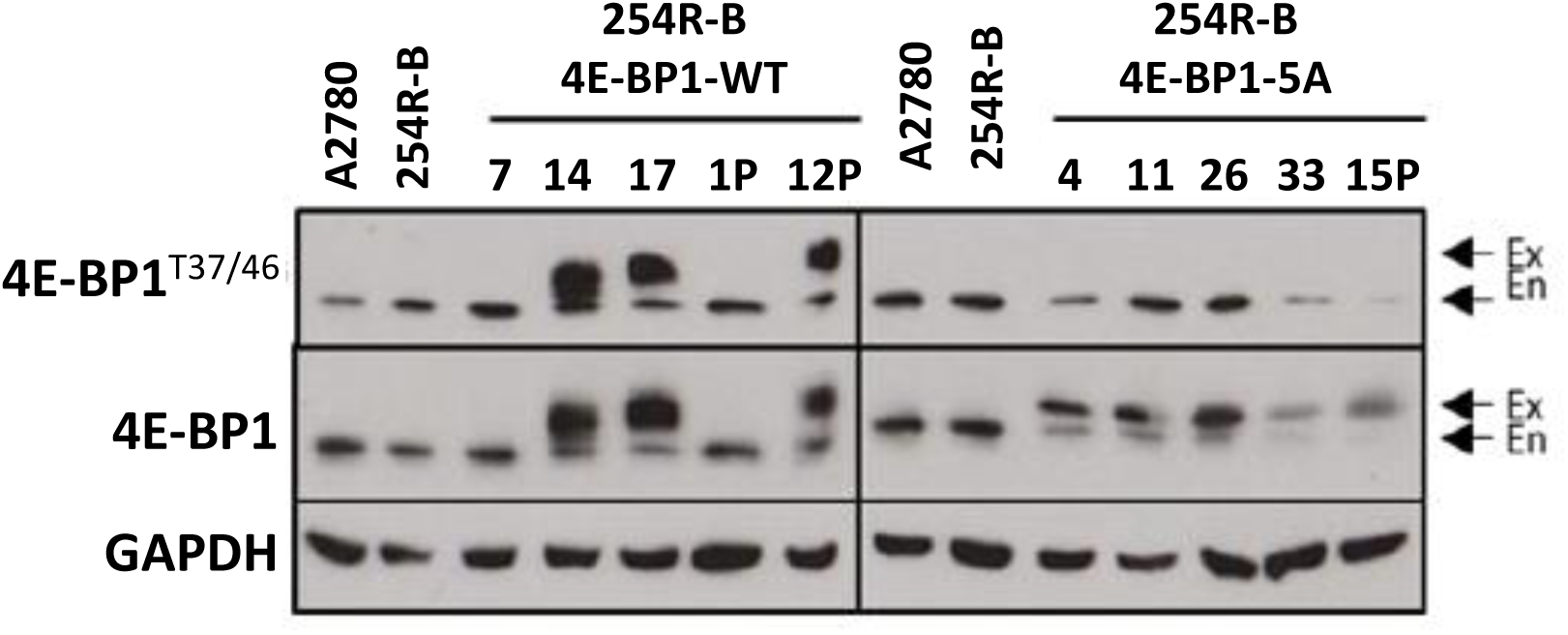
Expression of selected wild-type (WT) or 5A-mutant (5A) 4E-BP1 in selected stably transfected 254R-B cells. Western blot analysis of 4E-BP1 phosphorylation and expression in 254R-B pcDNA3-4E-BP1WT-HISMYC (WT)- or pcDNA3-4E-BP15A-HISMYC (5A)-transfected subclones with high 4E-BP1 expression. GAPDH was used as a loading control. Ex = exogenous band, En = endogenous band.

## Notes

### Competing Interest Statement

The authors have declared no competing interest.

## References

1. Glaviano, A., Foo, A.S.C., Lam, H.Y., Yap, K.C.H., Jacot, W., Jones, R.H., Eng, H., Nair, M.G., Makvandi, P., Geoerger, B., et al. (2023). PI3K/AKT/mTOR signaling transduction pathway and targeted therapies in cancer. Mol Cancer 22, 138. 10.1186/s12943-023-01827-6.

2. Kostaras, E., Kaserer, T., Lazaro, G., Heuss, S.F., Hussain, A., Casado, P., Hayes, A., Yandim, C., Palaskas, N., Yu, Y., et al. (2020). A systematic molecular and pharmacologic evaluation of AKT inhibitors reveals new insight into their biological activity. Br J Cancer 123, 542–555. 10.1038/s41416-020-0889-4.

3. Alves, C.L., and Ditzel, H.J. (2023). Drugging the PI3K/AKT/mTOR Pathway in ER+ Breast Cancer. IJMS 24, 4522. 10.3390/ijms24054522.

4. Davies, B.R., Greenwood, H., Dudley, P., Crafter, C., Yu, D.-H., Zhang, J., Li, J., Gao, B., Ji, Q., Maynard, J., et al. (2012). Preclinical Pharmacology of AZD5363, an Inhibitor of AKT: Pharmacodynamics, Antitumor Activity, and Correlation of Monotherapy Activity with Genetic Background. Molecular Cancer Therapeutics 11, 873–887. 10.1158/1535-7163.MCT-11-0824-T.

5. McHardy, T., Caldwell, J.J., Cheung, K.-M., Hunter, L.J., Taylor, K., Rowlands, M., Ruddle, R., Henley, A., De Haven Brandon, A., Valenti, M., et al. (2010). Discovery of 4-Amino-1-(7 *H* -pyrrolo[2,3-*d*]pyrimidin-4-yl)piperidine-4-carboxamides As Selective, Orally Active Inhibitors of Protein Kinase B (Akt). J. Med. Chem. 53, 2239–2249. 10.1021/jm901788j.

6. Addie, M., Ballard, P., Buttar, D., Crafter, C., Currie, G., Davies, B.R., Debreczeni, J., Dry, H., Dudley, P., Greenwood, R., et al. (2013). Discovery of 4-Amino-*N* -[(1 *S*)-1-(4-chlorophenyl)-3-hydroxypropyl]-1-(7 *H* -pyrrolo[2,3-*d*]pyrimidin-4-yl)piperidine-4-carboxamide (AZD5363), an Orally Bioavailable, Potent Inhibitor of Akt Kinases. J. Med. Chem. 56, 2059–2073. 10.1021/jm301762v.

7. Thomas, C., Lamoureux, F., Crafter, C., Davies, B.R., Beraldi, E., Fazli, L., Kim, S., Thaper, D., Gleave, M.E., and Zoubeidi, A. (2013). Synergistic Targeting of PI3K/AKT Pathway and Androgen Receptor Axis Significantly Delays Castration-Resistant Prostate Cancer Progression *In Vivo*. Molecular Cancer Therapeutics 12, 2342–2355. 10.1158/1535-7163.MCT-13-0032.

8. Li, J., Davies, B.R., Han, S., Zhou, M., Bai, Y., Zhang, J., Xu, Y., Tang, L., Wang, H., Liu, Y.J., et al. (2013). The AKT inhibitor AZD5363 is selectively active in PI3KCA mutant gastric cancer, and sensitizes a patient-derived gastric cancer xenograft model with PTEN loss to Taxotere. J Transl Med 11, 241. 10.1186/1479-5876-11-241.

9. Ribas, R., Pancholi, S., Guest, S.K., Marangoni, E., Gao, Q., Thuleau, A., Simigdala, N., Polanska, U.M., Campbell, H., Rani, A., et al. (2015). AKT Antagonist AZD5363 Influences Estrogen Receptor Function in Endocrine-Resistant Breast Cancer and Synergizes with Fulvestrant (ICI182780) *In Vivo*. Molecular Cancer Therapeutics 14, 2035–2048. 10.1158/1535-7163.MCT-15-0143.

10. Davies, B.R., Guan, N., Logie, A., Crafter, C., Hanson, L., Jacobs, V., James, N., Dudley, P., Jacques, K., Ladd, B., et al. (2015). Tumors with AKT1E17K Mutations Are Rational Targets for Single Agent or Combination Therapy with AKT Inhibitors. Molecular Cancer Therapeutics 14, 2441–2451. 10.1158/1535-7163.MCT-15-0230.

11. Gris-Oliver, A., Palafox, M., Monserrat, L., Brasó-Maristany, F., Òdena, A., Sánchez-Guixé, M., Ibrahim, Y.H., Villacampa, G., Grueso, J., Parés, M., et al. (2020). Genetic Alterations in the PI3K/AKT Pathway and Baseline AKT Activity Define AKT Inhibitor Sensitivity in Breast Cancer Patient-derived Xenografts. Clinical Cancer Research 26, 3720–3731. 10.1158/1078-0432.CCR-19-3324.

12. Banerji, U., Dean, E.J., Pérez-Fidalgo, J.A., Batist, G., Bedard, P.L., You, B., Westin, S.N., Kabos, P., Garrett, M.D., Tall, M., et al. (2018). A Phase I Open-Label Study to Identify a Dosing Regimen of the Pan-AKT Inhibitor AZD5363 for Evaluation in Solid Tumors and in *PIK3CA* -Mutated Breast and Gynecologic Cancers. Clinical Cancer Research 24, 2050–2059. 10.1158/1078-0432.CCR-17-2260.

13. Jones, R.H., Casbard, A., Carucci, M., Cox, C., Butler, R., Alchami, F., Madden, T.-A., Bale, C., Bezecny, P., Joffe, J., et al. (2020). Fulvestrant plus capivasertib versus placebo after relapse or progression on an aromatase inhibitor in metastatic, oestrogen receptor-positive breast cancer (FAKTION): a multicentre, randomised, controlled, phase 2 trial. The Lancet Oncology 21, 345–357. 10.1016/S1470-2045(19)30817-4.

14. Qureshi, Z., Altaf, F., Khanzada, M., Zaheer, Z., Fatima, E., and Bakhtiar, M. (2024). Capivasertib in Hormone Receptor-Positive, Human Epidermal Growth Factor Receptor 2-Negative advanced breast cancer. Current Problems in Cancer 51, 101114. 10.1016/j.currproblcancer.2024.101114.

15. Dilawari, A., Buturla, J., Osgood, C., Gao, X., Chen, W., Ricks, T.K., Schaefer, T., Avasarala, S., Reyes Turcu, F., Pathak, A., et al. (2024). US Food and Drug Administration Approval Summary: Capivasertib With Fulvestrant for Hormone Receptor–Positive, Human Epidermal Growth Factor Receptor 2–Negative Locally Advanced or Metastatic Breast Cancer With PIK3CA / AKT1 / PTEN Alterations. JCO 42, 4103–4113. 10.1200/JCO.24.00427.

16. De, S.K. (2025). Capivasertib: First Approved AKT inhibitor for the Treatment of Patients with Breast Cancer. ACAMC 25, 371–377. 10.2174/0118715206360571241126080725.

17. Oliveira, M., Rugo, H.S., Howell, S.J., Dalenc, F., Cortes, J., Gomez, H.L., Hu, X., Toi, M., Jhaveri, K., Krivorotko, P., et al. (2024). Capivasertib and fulvestrant for patients with hormone receptor-positive, HER2-negative advanced breast cancer (CAPItello-291): patient-reported outcomes from a phase 3, randomised, double-blind, placebo-controlled trial. The Lancet Oncology 25, 1231–1244. 10.1016/S1470-2045(24)00373-5.

18. Stubbs, N.M., Roady, T.S., Schwermann, M.P., Eteshola, E.O., MacDonald, W.J., Purcell, C., Ryspayeva, D., Verovkina, N., Tajiknia, V., Ghandali, M., et al. (2025). Acquired resistance to molecularly targeted therapies for cancer. Cancer Drug Resist. 10.20517/cdr.2024.189.

19. Zhang, H., Jiang, R., Zhu, J., Sun, K., Huang, Y., Zhou, H., Zheng, Y., and Wang, X. (2024). PI3K/AKT/mTOR signaling pathway: an important driver and therapeutic target in triple-negative breast cancer. Breast Cancer 31, 539–551. 10.1007/s12282-024-01567-5.

20. Browne, I.M., and Okines, A.F.C. (2024). Resistance to Targeted Inhibitors of the PI3K/AKT/mTOR Pathway in Advanced Oestrogen-Receptor-Positive Breast Cancer. Cancers 16, 2259. 10.3390/cancers16122259.

21. Dunn, S., Eberlein, C., Yu, J., Gris-Oliver, A., Ong, S.H., Yelland, U., Cureton, N., Staniszewska, A., McEwen, R., Fox, M., et al. (2022). AKT-mTORC1 reactivation is the dominant resistance driver for PI3Kβ/AKT inhibitors in PTEN-null breast cancer and can be overcome by combining with Mcl-1 inhibitors. Oncogene 41, 5046–5060. 10.1038/s41388-022-02482-9.

22. Ediriweera, M.K., Tennekoon, K.H., and Samarakoon, S.R. (2019). Role of the PI3K/AKT/mTOR signaling pathway in ovarian cancer: Biological and therapeutic significance. Seminars in Cancer Biology 59, 147–160. 10.1016/j.semcancer.2019.05.012.

23. Mabuchi, S., Kuroda, H., Takahashi, R., and Sasano, T. (2015). The PI3K/AKT/mTOR pathway as a therapeutic target in ovarian cancer. Gynecologic Oncology 137, 173–179. 10.1016/j.ygyno.2015.02.003.

24. Li, H., Zeng, J., and Shen, K. (2014). PI3K/AKT/mTOR signaling pathway as a therapeutic target for ovarian cancer. Arch Gynecol Obstet 290, 1067–1078. 10.1007/s00404-014-3377-3.

25. Gasparri, M., Bardhi, E., Ruscito, I., Papadia, A., Farooqi, A., Marchetti, C., Bogani, G., Ceccacci, I., Mueller, M., and Benedetti Panici, P. (2017). PI3K/AKT/mTOR Pathway in Ovarian Cancer Treatment: Are We on the Right Track? Geburtshilfe Frauenheilkd 77, 1095–1103. 10.1055/s-0043-118907.

26. Dobbin, Z., and Landen, C. (2013). The Importance of the PI3K/AKT/MTOR Pathway in the Progression of Ovarian Cancer. IJMS 14, 8213–8227. 10.3390/ijms14048213.

27. Van Der Ploeg, P., Uittenboogaard, A., Thijs, A.M.J., Westgeest, H.M., Boere, I.A., Lambrechts, S., Van De Stolpe, A., Bekkers, R.L.M., and Piek, J.M.J. (2021). The effectiveness of monotherapy with PI3K/AKT/mTOR pathway inhibitors in ovarian cancer: A meta-analysis. Gynecologic Oncology 163, 433–444. 10.1016/j.ygyno.2021.07.008.

28. Michaelis, M., Wass, M.N., and Cinatl, J. (2019). Drug-adapted cancer cell lines as preclinical models of acquired resistance. CDR. 10.20517/cdr.2019.005.

29. Grimsley, H.E., Antczak, M., Reddin, I.G., Weiler, N., McLaughlin, K.-M., Rothweiler, F., Haas, J., Nist, A., Mernberger, M., Stiewe, T., et al. (2025). Using a novel panel of drug-resistant triple-negative breast cancer cell lines to identify candidate therapeutic targets and biomarkers. Cancer Letters 624, 217754. 10.1016/j.canlet.2025.217754.

30. Zuccotti, P., and Modelska, A. (2016). Studying the Translatome with Polysome Profiling. In Post-Transcriptional Gene Regulation Methods in Molecular Biology., E. Dassi, ed. (Springer New York), pp. 59–69. 10.1007/978-1-4939-3067-8_4.

31. Beaufort, C.M., Helmijr, J.C.A., Piskorz, A.M., Hoogstraat, M., Ruigrok-Ritstier, K., Besselink, N., Murtaza, M., Van IJcken, W.F.J., Heine, A.A.J., Smid, M., et al. (2014). Ovarian Cancer Cell Line Panel (OCCP): Clinical Importance of In Vitro Morphological Subtypes. PLoS ONE 9, e103988. 10.1371/journal.pone.0103988.

32. Oda, K., Okada, J., Timmerman, L., Rodriguez-Viciana, P., Stokoe, D., Shoji, K., Taketani, Y., Kuramoto, H., Knight, Z.A., Shokat, K.M., et al. (2008). PIK3CA Cooperates with Other Phosphatidylinositol 3′-Kinase Pathway Mutations to Effect Oncogenic Transformation. Cancer Research 68, 8127–8136. 10.1158/0008-5472.CAN-08-0755.

33. O’Reilly, K.E., Rojo, F., She, Q.-B., Solit, D., Mills, G.B., Smith, D., Lane, H., Hofmann, F., Hicklin, D.J., Ludwig, D.L., et al. (2006). mTOR Inhibition Induces Upstream Receptor Tyrosine Kinase Signaling and Activates Akt. Cancer Research 66, 1500–1508. 10.1158/0008-5472.CAN-05-2925.

34. Breuleux, M., Klopfenstein, M., Stephan, C., Doughty, C.A., Barys, L., Maira, S.-M., Kwiatkowski, D., and Lane, H.A. (2009). Increased AKT S473 phosphorylation after mTORC1 inhibition is rictor dependent and does not predict tumor cell response to PI3K/mTOR inhibition. Molecular Cancer Therapeutics 8, 742–753. 10.1158/1535-7163.MCT-08-0668.

35. Thoreen, C.C., Kang, S.A., Chang, J.W., Liu, Q., Zhang, J., Gao, Y., Reichling, L.J., Sim, T., Sabatini, D.M., and Gray, N.S. (2009). An ATP-competitive Mammalian Target of Rapamycin Inhibitor Reveals Rapamycin-resistant Functions of mTORC1. Journal of Biological Chemistry 284, 8023–8032. 10.1074/jbc.M900301200.

36. Bitterman, P.B., and Polunovsky, V.A. (2015). eIF4E-mediated translational control of cancer incidence. Biochimica et Biophysica Acta (BBA) - Gene Regulatory Mechanisms 1849, 774–780. 10.1016/j.bbagrm.2014.09.007.

37. Nandagopal, N., and Roux, P.P. (2015). Regulation of global and specific mRNA translation by the mTOR signaling pathway. Translation 3, e983402. 10.4161/21690731.2014.983402.

38. Qin, X., Jiang, B., and Zhang, Y. (2016). 4E-BP1, a multifactor regulated multifunctional protein. Cell Cycle 15, 781–786. 10.1080/15384101.2016.1151581.

39. Roux, P.P., and Topisirovic, I. (2012). Regulation of mRNA Translation by Signaling Pathways. Cold Spring Harbor Perspectives in Biology 4, a012252– a012252. 10.1101/cshperspect.a012252.

40. Leppek, K., Das, R., and Barna, M. (2018). Functional 5′ UTR mRNA structures in eukaryotic translation regulation and how to find them. Nat Rev Mol Cell Biol 19, 158–174. 10.1038/nrm.2017.103.

41. Herbert, T.P., Tee, A.R., and Proud, C.G. (2002). The Extracellular Signal-regulated Kinase Pathway Regulates the Phosphorylation of 4E-BP1 at Multiple Sites. Journal of Biological Chemistry 277, 11591–11596. 10.1074/jbc.M110367200.

42. Wang, X., Li, W., Parra, J.-L., Beugnet, A., and Proud, C.G. (2003). The C Terminus of Initiation Factor 4E-Binding Protein 1 Contains Multiple Regulatory Features That Influence Its Function and Phosphorylation. Molecular and Cellular Biology 23, 1546–1557. 10.1128/MCB.23.5.1546-1557.2003.

43. André, F., Ciruelos, E., Rubovszky, G., Campone, M., Loibl, S., Rugo, H.S., Iwata, H., Conte, P., Mayer, I.A., Kaufman, B., et al. (2019). Alpelisib for *PIK3CA* - Mutated, Hormone Receptor–Positive Advanced Breast Cancer. N Engl J Med 380, 1929–1940. 10.1056/NEJMoa1813904.

44. Savill, K.M.Z., Lee, B.B., Oeh, J., Lin, J., Lin, E., Chung, W.-J., Young, A., Chen, W., Miś, M., Mesh, K., et al. (2022). Distinct resistance mechanisms arise to allosteric vs. ATP-competitive AKT inhibitors. Nat Commun 13, 2057. 10.1038/s41467-022-29655-0.

45. Stanevich, V., Jiang, L., Satyshur, K.A., Li, Y., Jeffrey, P.D., Li, Z., Menden, P., Semmelhack, M.F., and Xing, Y. (2011). The Structural Basis for Tight Control of PP2A Methylation and Function by LCMT-1. Molecular Cell 41, 331–342. 10.1016/j.molcel.2010.12.030.

46. Yang, X., Zhong, W., and Cao, R. (2020). Phosphorylation of the mRNA cap-binding protein eIF4E and cancer. Cellular Signalling 73, 109689. 10.1016/j.cellsig.2020.109689.

47. Hong, D.S., Kurzrock, R., Oh, Y., Wheler, J., Naing, A., Brail, L., Callies, S., André, V., Kadam, S.K., Nasir, A., et al. (2011). A Phase 1 Dose Escalation, Pharmacokinetic, and Pharmacodynamic Evaluation of eIF-4E Antisense Oligonucleotide LY2275796 in Patients with Advanced Cancer. Clinical Cancer Research 17, 6582–6591. 10.1158/1078-0432.CCR-11-0430.

48. Borden, K.L.B., and Culjkovic-Kraljacic, B. (2010). Ribavirin as an anti-cancer therapy: acute myeloid leukemia and beyond? Leukemia & Lymphoma 51, 1805– 1815. 10.3109/10428194.2010.496506.

49. Shi, F., Len, Y., Gong, Y., Shi, R., Yang, X., Naren, D., and Yan, T. (2015). Ribavirin Inhibits the Activity of mTOR/eIF4E, ERK/Mnk1/eIF4E Signaling Pathway and Synergizes with Tyrosine Kinase Inhibitor Imatinib to Impair Bcr-Abl Mediated Proliferation and Apoptosis in Ph+ Leukemia. PLoS ONE 10, e0136746. 10.1371/journal.pone.0136746.

50. Assouline, S., Culjkovic, B., Cocolakis, E., Rousseau, C., Beslu, N., Amri, A., Caplan, S., Leber, B., Roy, D.-C., Miller, W.H., et al. (2009). Molecular targeting of the oncogene eIF4E in acute myeloid leukemia (AML): a proof-of-principle clinical trial with ribavirin. Blood 114, 257–260. 10.1182/blood-2009-02-205153.

51. Smith, K.A., Zhou, B., Avdulov, S., Benyumov, A., Peterson, M., Liu, Y., Okon, A., Hergert, P., Braziunas, J., Wagner, C.R., et al. (2015). Transforming Growth Factor-β1 Induced Epithelial Mesenchymal Transition is blocked by a chemical antagonist of translation factor eIF4E. Sci Rep 5, 18233. 10.1038/srep18233.

52. Chen, E.Z., Jacobson, B.A., Patel, M.R., Okon, A.M., Li, S., Xiong, K., Vaidya, A.J., Bitterman, P.B., Wagner, C.R., and Kratzke, R.A. (2014). Small-molecule inhibition of oncogenic eukaryotic protein translation in mesothelioma cells. Invest New Drugs 32, 598–603. 10.1007/s10637-014-0076-7.

53. De, A., Jacobson, B.A., Peterson, M.S., Jay-Dixon, J., Kratzke, M.G., Sadiq, A.A., Patel, M.R., and Kratzke, R.A. (2018). 4EGI-1 represses cap-dependent translation and regulates genome-wide translation in malignant pleural mesothelioma. Invest New Drugs 36, 217–229. 10.1007/s10637-017-0535-z.

54. Descamps, G., Gomez-Bougie, P., Tamburini, J., Green, A., Bouscary, D., Maïga, S., Moreau, P., Le Gouill, S., Pellat-Deceunynck, C., and Amiot, M. (2012). The cap-translation inhibitor 4EGI-1 induces apoptosis in multiple myeloma through Noxa induction. Br J Cancer 106, 1660–1667. 10.1038/bjc.2012.139.

55. Wang, H., Huang, F., Wang, J., Wang, P., Lv, W., Hong, L., Li, S., and Zhou, J. (2015). The synergistic inhibition of breast cancer proliferation by combined treatment with 4EGI-1 and MK2206. Cell Cycle 14, 232–242. 10.4161/15384101.2014.977096.

56. Roux, P.P., and Topisirovic, I. (2012). Regulation of mRNA Translation by Signaling Pathways. Cold Spring Harbor Perspectives in Biology 4, a012252– a012252. 10.1101/cshperspect.a012252.

